# Aridity Drives Climatic Specialization and Phylogenetic Clustering in Terrestrial Vertebrates

**DOI:** 10.1101/2025.10.13.681566

**Authors:** Héctor Tejero-Cicuéndez, Gabriel Arellano, Iris Menéndez, Joan Garcia-Porta

**Author notes:** Corresponding authors. /.

## Abstract

Arid environments constitute one of the main terrestrial biomes on Earth and are predicted to expand under ongoing climate change. Studying the effect of arid conditions on biodiversity patterns and ecological dynamics is therefore critical to understand the evolutionary fate of desert-adapted faunas. Here we provide a global assessment of how aridity influences climatic niche breadth and phylogenetic structure across terrestrial vertebrates. Using distributional data for over 34,000 species of amphibians, squamates, birds, and mammals, combined with environmental data and nearly complete phylogenies of these clades, we quantified species’ positions along aridity gradients and their climatic niche breadth, and evaluated phylogenetic clustering at multiple spatial scales. Across all groups, species occupying more arid environments exhibited significantly narrower climatic niches than those from non-arid habitats, supporting the hypothesis that ecological filtering in deserts promotes climatic specialization. Further, analyses of system-wide community phylogenetic structure revealed significant phylogenetic clustering in most arid systems for all clades except birds, underscoring that deserts broadly act as ecological filters at large scale, but this effect is subjected to clade-specific traits. However, finer-scale analyses showed significant phylogenetic overdispersion in several cases, suggesting that competition and micro-niche dynamics might be stronger determinants of phylogenetic structure at the local-to-regional level. Together, our results demonstrate that drylands act as ecological filters shaping both functional and evolutionary dimensions of biodiversity, with implications for predicting biotic responses to desertification and climate change.

## Introduction

Arid and semiarid environments cover approximately 40% of the planet’s land surface (Gaur and Squires 2018). With climate change driving the continued expansion of drylands (Mirzabaev et al. 2022), it is increasingly crucial to explore how organisms colonize and adapt in such conditions.

Arid regions are characterized by low water availability and, very often, by extreme temperatures. Therefore, they likely demand specialized traits for thermoregulation and water conservation. As a result, species in these environments often exhibit unique physiological and morphological traits oriented to cope with such extreme conditions (Bentley 1966; Kronfeld and Shkolnik 1996; Williams and Tieleman 2002; Osborne et al. 2020; Chabaud et al. 2022). Such specialization, however, may limit performance in less extreme environments, constraining the climatic niche breadth of arid-adapted species relative to those in non-arid regions (Moreno-Rueda 2014; Alhajeri and Steppan 2018).

Alternatively, some organisms may cope with arid conditions through phenotypic or behavioral plasticity rather than relying solely on evolved physiological or morphological adaptations (Muñoz et al. 2016; Muñoz 2022). Behavioral flexibility can lessen the need for extensive physiological or morphological changes, for example by shifting activity to cooler periods of the day (Kearney et al. 2009; Rafiq et al. 2023) or selecting favorable microhabitats such as burrows (Goller et al. 2014) (Figure 1). Another possibility is that organisms colonize arid regions by tracking their preferred niche, even if it is rare within these environments (e. g., oases or riparian zones). Such strategies would reduce differences in climatic niche breadth between arid and non-arid species.

**Figure 1.**
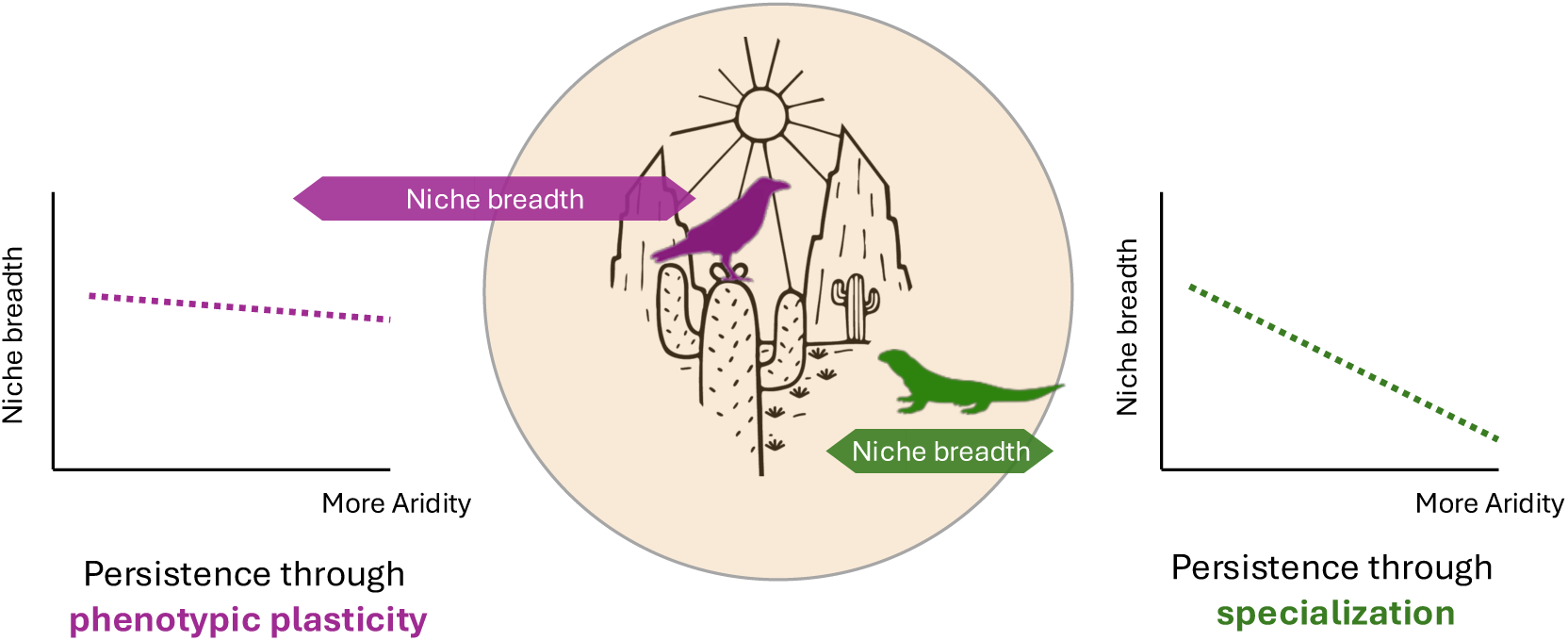
Conceptual representation of the two hypotheses that potentially explain the colonization and persistence in arid regions: phenotypic plasticity (left), which results in arid species having similar niche breadths to those of non-arid species, and climatic specialization (right), which is reflected in the narrower niche breadths of arid species relative to non-arid taxa.

These alternative strategies have important consequences for the expected strength of ecological filtering occurring in these environments. If successful colonization of arid regions generally relies on specialized traits (which may be evolutionarily costly to develop), high ecological filtering is expected, leading to communities dominated by functionally similar and closely related species (e.g., Gong et al. 2019; Doby et al. 2022). Conversely, if coping with arid conditions occurs through phenotypic plasticity or niche tracking, lower levels of ecological filtering are expected, generating functionally and phylogenetically diverse assemblages within arid habitats.

The relative contribution of each scenario is likely shaped by a scale-dependent combination of organismal and environmental factors. From an organismal perspective, different taxonomic groups may vary in their capacity for phenotypic plasticity or their evolutionary potential to develop traits suited to arid conditions, potentially leading to different strategies for persistence in such environments (McKechnie and Wolf 2004; Baldwin et al. 2022). From an environmental perspective, arid systems – despite their broad climatic similarity – exhibit considerable ecological and environmental heterogeneity. For instance, some deserts may impose stronger selective pressures for specialization due to greater environmental harshness. Moreover, even under similar levels of harshness, variation in the age of desert systems provides differing amounts of evolutionary time for climatic specialization to develop, further contributing to differences in the characteristics of species assemblages across arid regions (Tejero-Cicuéndez et al. 2022). All this considered, different geographic scales can also theoretically produce multiple signals of phylogenetic clustering in arid regions. At smaller spatial scales, where environments tend to be more uniform and competition is stronger, niche segregation can occur only at the microniche level. In such cases, we expect narrower climatic niches coupled with low phylogenetic clustering, as competition favors ecological differentiation. At larger scales, however, greater environmental heterogeneity can accommodate more closely related species, thereby increasing phylogenetic clustering (Webb et al. 2002; Cavender-Bares et al. 2009; Park et al. 2020).

In this study, we evaluate how these factors interact to shape climatic niche breadth and phylogenetic diversity in drylands worldwide. Specifically, we test whether species in arid environments exhibit narrower climatic niches and stronger phylogenetic clustering than those in non-arid regions, and whether these patterns vary among clades, across desert systems, and between spatial scales. By integrating species-level climatic data with community phylogenetic analyses, our study provides a global perspective on how drylands act as drivers of specialization and ecological filtering in terrestrial vertebrates.

## Materials & Methods

### Characterization of the climatic niches of tetrapod species

#### Distributional data

As a first step, we characterized the position and breadth of the climatic niches of tetrapod species by intersecting their distributional ranges with climatic data. Distributional ranges were sourced from three main databases: For amphibians and mammals, we downloaded shapefiles from the International Union for Conservation of Nature (IUCN 2022), while data for birds were acquired from BirdLife International (https://datazone.birdlife.org/) and reptile range maps were sourced from (Roll et al. 2021). Each dataset was filtered to retain only extant and native species; for migratory birds, we included only areas classified as “resident” or “breeding” in the BirdLife data. The final datasets included 7,175 species of amphibians, 10,585 species of squamates, 10,814 species of birds, and 5,599 species of mammals. See Appendix 1 for the R code used to filter the bird distribution layers.

#### Climatic data

We used three sources of global climatic and environmental data. First, we used the WorldClim dataset v2.1 (Fick and Hijmans 2017) downloaded at a 10-minute spatial resolution (∼18.5 km). This dataset contains high-resolution global climate data for the period 1970-2000, which includes 19 bioclimatic variables that are relevant for living taxa. Secondly, we used the Global Aridity Index (v3, Zomer et al. 2022), a spatial raster layer that quantifies an aridity index (AI) estimated from precipitation and evapotranspiration values (Greve et al. 2019). Low AI values (down to zero) correspond to high-aridity regions, while high values represent humid regions. The aridity raster data, with a spatial resolution of approximately 1 km, was resampled to align with the resolution of the WorldClim dataset (∼18.5 km). Finally, we used the Biome 13 “Desert and xeric shrublands” defined by Olson et al. (2001), to identify the main arid systems in the world. This biome delimits several arid ecoregions that can be lumped into eight main arid zones characterized by significant arid and hyper-arid conditions: Atacama, Australian desert, Central Asian desert, Gobi, Kalahari, North American desert, Persian desert, and Saharo-Arabian desert (Tejero-Cicuéndez et al. 2022).

### Characterization of aridity niche and climatic niche breadth

The first step is to position each species along the aridity gradient through multiple approaches. We overlapped the distribution of each species with the AI raster, which allowed us to generate three complementary variables reflecting the association between species and aridity: (i) considering cells with AI values < 0.25 as arid (see Supplementary Figure 1), we generated a continuous variable by estimating the percentage of arid cells within the distribution of each species; (ii) we generated a categorical variable applying a threshold of 50% of arid cells within each species’ distribution to categorize them as arid (≥ 50%) and non-arid (< 50%); (iii) we calculated the median AI value for each species to position them along a continuous aridity gradient.

In addition to this, we assigned species to the main desert systems (see above). This approach closely aligns with the aridity threshold used to define arid and non-arid species (see Supplementary Figure 1). A species was categorized as belonging to a particular desert if at least 15% of its distribution overlapped with the desert or if it occupied at least 15% of the desert’s total area (Appendix 2 shows how the species numbers in each desert change with different thresholds; Tejero-Cicuéndez et al. 2022).

To estimate the climatic niche breadth for each species, we conducted a Principal Component Analysis (PCA) using all bioclimatic variables from the WorldClim dataset. For each species, we then calculated, based on the overlapping raster cells with the species’ distribution area, the range encompassing the central 95% of the variation along the first two principal components (PCs).

### Phylogenetic data

To explore the relationship between niche breadth and aridity while accounting for the non-independence of species due to evolutionary relatedness, and to compare phylogenetic clustering between communities in arid and non-arid environments, we used nearly complete, time-calibrated phylogenetic trees for each major vertebrate group. These included the trees from Jetz and Pyron (2018) for amphibians, Tonini et al. (2016) for squamates, Jetz et al. (2012) for birds, and Upham et al. (2019) for mammals. These phylogenies were generated through an initial estimation of topology and divergence times based on available genetic data, followed by the imputation of species lacking genetic data using a stochastic polytomy resolver (Thomas et al. 2013).

### Exploring per-species global relationship between niche breadth and aridity

To investigate how climatic specialization (understood here by narrowness of niche breadth) varies across the aridity gradient and vertebrate groups, we employed three complementary approaches.

#### Phylogenetic Analyses of Variance

We first visualized the differences in climatic niche breadth between the species categorized as arid (species in which arid cells constituted 50% or more of their distribution) and non-arid in each vertebrate group, by means of violin plots. We tested significance using a phylogenetic analyses of variance (ANOVA) as implemented in the R package RRPP (v2.1.2; Collyer and Adams 2018, 2025).

#### Phylogenetic regression models

Secondly, we conducted phylogenetic linear regression models to examine the covariation between niche breadth, defined as the central 95% of variation along the first two principal components, and aridity across terrestrial vertebrate species. To this end, we used the continuous variables described above to characterize the species’ aridity niche: (1) the median position of each species along the aridity gradient, and (2) the percentage of arid grid cells within each species’ distribution. Because the evolutionary model assumed in the regression can substantially influence its outcome (Freckleton and Harvey 2006), we compared multiple models and transformations available in the R package phylolm (Ho and Ané 2014): Brownian Motion, Ornstein-Uhlenbeck with random root, Ornstein-Uhlenbeck with fixed root, lambda transformation, and Early Burst. For each analysis, we selected the model with the lowest Akaike Information Criterion (AIC; Akaike 1973).

Given the strong correlation between range size and niche breadth (Slatyer et al. 2013), range size may act as a confounding factor: species in arid environments could appear to have narrower niches simply due to more restricted distributions. To control for this, we conducted an additional phylogenetic multiple linear regression including log-transformed range size as a covariate.

To further disentangle the effect of range size, we applied an alternative approach. We first regressed niche breadth against range size using a local regression (LOESS) model with parameters α = 0.75 and degree = 1 and then extracted the residuals. These residuals, representing niche breadth corrected for range size, were then used in a subsequent phylogenetic linear regression with phylolm to reassess covariation with aridity.

### Exploring the relationship between niche breadth and aridity among desert systems

Thirdly, we explored how the degree of climatic specialization differs across arid systems, examining the relationship between niche breadth and species’ median aridity for each arid system. This allowed us to identify region-specific differences in the degree of specialization along the aridity gradient, which may arise due to differences among deserts resulting in ecological pressures of varying strength (system-specific environmental filtering). To do this, we inspected the slope of a phylogenetic regression model setting as predictor term the interaction between aridity and the presence/absence of each species in each arid system.

However, instead of selecting only species within the arid systems (which would only help us explore differences among arid-adapted species), we also included species living in the surrounding non-arid areas to each desert. This allows us to explore system-specific niche specialization along the aridity gradient including species from arid and non-arid environments, while controlling for potential eco-evolutionary differences of faunas inhabiting different landmasses or distant regions. With that purpose, we defined eight buffer regions, each containing one desert and its surrounding area within 15 degrees latitude and longitude from the desert’s boundaries (see Supplementary Figure 2). The presence/absence of each species in each of these buffer regions was then added as an interaction term in a general phylogenetic regression model (including all regions), and we additionally performed individual phylogenetic regression models for each buffer region and compared the slopes. For both approaches we used phylolm (Ho and Ané 2014).

### Analyses of phylogenetic clustering

Life in arid environments often requires special adaptations for water balance and temperature regulation, which can be challenging to evolve. This may lead to strong habitat filtering, increasing the phylogenetic clustering of arid-adapted communities (stress-dominance hypothesis; Patrick and Stevens 2016; Ramm et al. 2018). Alternatively, if arid environments are colonized through phenotypic plasticity—achieved via multiple evolutionary routes—a weaker habitat filtering is expected, leading to a low phylogenetic clustering in desert communities. To evaluate these scenarios, we compared the observed phylogenetic clustering of species assemblages in arid environments with the expected phylogenetic clustering based on a null model.

Since the ecological processes influencing patterns of phylogenetic diversity (e.g., ecological filtering or biotic interactions) might be scale-dependent (e.g., Patrick and Stevens 2016), we performed this comparison at two geographic scales. At a large scale, we focused on major desert systems, quantifying different phylogenetic metrics in each system as a whole and comparing them with a null model consisting of 100 simulated species assemblages. Given the potentially different evolutionary history of species in different landmasses (e.g., dispersal limitations likely make species assemblages in North American more different to those in Africa than to others in North America), we built the null species assemblages using the eight buffer regions defined above (see Supplementary Figure 2). Phylogenetic clustering in each arid system was then compared to that of the 100 species assemblages composed of randomly selected species from the buffer regions, keeping the number of species equal to that of the arid system (Supplementary Figure 3). This allows us to evaluate the potential role of the desert as an ecological filter, by testing whether arid systems harbor particularly high or low levels of phylogenetic clustering. For each group of terrestrial vertebrates, phylogenetic trees were pruned to include only species found within each assemblage. To quantify phylogenetic clustering, we calculated three metrics: phylogenetic diversity (PD; Faith 1992), mean phylogenetic pairwise distance (MPD), and mean nearest taxon distance (MNTD). All these metrics provide information about phylogenetic structure, but they do so in different manners: i) PD is defined as the sum of the lengths of all the branches on the tree that span the species of the set (Faith 1992); ii) MPD is the mean phylogenetic distance (i.e., branch length) among all pairs of species within a community (Tsirogiannis and Sandel 2014); iii) MNTD, also known as mean nearest neighbor distance (MNND), is the mean distance between each species within a community and its closest relative (Webb et al. 2002). We further computed standardized effect sizes (SES) for these metrics, which measure the difference between the observed phylogenetic structure and the mean expected under null models, standardized by the variance of expected values (Gotelli and Rohde 2002). SES values were calculated using the *ses.pd*, *ses.mpd*, and *ses.mntd* functions in the R package picante (Kembel et al. 2010), with 999 randomizations.

At a finer scale, we analyzed phylogenetic clustering of species assemblages at a resolution of 270 × 270 km. In this case, we generated a map of residual PD (residuals of a LOESS regression of PD on species richness) and calculated the observed average residual PD in the 25% most arid locations. The 25% threshold was selected over several others (10, 15, 20, 30, 35, 40, 45) after corroborating that it resulted in an adequate representation of arid regions worldwide (Supplementary Figure 4). We did this for the 25% most arid locations overall, and also separating locations by their corresponding arid system, to gain a more nuanced perspective of the underlying phylogenetic diversity dynamics among deserts at a regional scale. Then, we tested whether this observed residual PD in the most arid sites was significantly lower or higher than the average obtained from 999 datasets simulated using a null model. Our novel null model consisted of random rotations of aridity values across the Earth’s surface, but constrained to the land surface only to avoid the useless expectation of many species present in oceans. Our procedure was as follows:

1. We simulated an “only-land” sphere by matching uniformly distributed locations within the continents to uniformly distributed locations in a sphere, in a one-to-one assignment. This effectively stretched all continents or land masses as if they covered the whole Earth (Supplementary Figure 5). This was performed by using the algorithm of Jonker and Volgenant (1987) (a faster alternative to the Hungarian algorithm; Munkres 1957) as implemented in the R package TreeDist (Smith 2020).
2. Once each single location within land surface was assigned to one of the locations within the “only-land” sphere, we proceeded to rotate these 3D locations in a random angle, with an after-rotation adjustment so that the new (null) coordinates of each location matched exactly the coordinates of one of the original locations, using a fast k-nearest neighbor searching algorithm implemented in the R package ‘FNN’ (Beygelzimer et al. 2024). Moreover, after each random rotation, we implemented an optimization step to minimize the error between the original autocorrelation of the environmental variable and the autocorrelation of the rotated variable.
3. After rotating in the “only-land” sphere, we translocated each rotated point to the corresponding coordinates in the Earth, so that each one of the original sites is matched with a null location within land.

These random rotations break any spatial association between environmental variables and species geographic patterns, while the spatial structure of both the environment and the species distributions is respected. Therefore, it constitutes a better null model for exploring the extent to which two sets of geographically distributed variables are related than other null-model approaches such as random sampling of geographic sites.

Doing this procedure 999 times, we ended up with 999 “null environments”, in which the environmental variables (in our case, the aridity values) are rotated in different angles and keep most of the autocorrelation and spatial structure of the original variable (Supplementary Figure 6). We assessed the significance of the observed residual PD in the 25% most arid locations by comparing it to the average residual PD in the 25% most arid locations after each null rotation.

## Results

### Characterization of the climatic niche

Our method for identifying species distribution along the aridity gradients of Earth confirms the expected notion that arid environments are generally less diverse than non-arid environments. Specifically, for amphibians, 3.7% of species were associated with arid environments; for reptiles, 24%; for birds, 10.7%; and for mammals, 14.7% (Supplementary Figure 7).

Principal Component Analysis (PCA) conducted on the WorldClim dataset captured 74% of the total variance in climatic data with the first two principal components (PCs): PC1 accounted for 56% of the variance, and PC2 for 18% (Supplementary Figure 8). PC1 was dominated by temperature-related variables, including annual mean temperature, minimum temperature of the coldest month, and minimum temperatures of the driest, warmest, and coldest quarters. In contrast, PC2 represented a mix of temperature- and precipitation-related variables, such as mean diurnal range, maximum temperature of the warmest month, and precipitation of the driest month, driest quarter, and coldest quarter (Supplementary Figure 9).

Given the high variance explained by PC1, we used it as our primary proxy for climatic niche breadth in subsequent analyses. However, all analyses were repeated using PC2, consistently yielding results that aligned with those derived from PC1 (see Supplementary).

### Relationship between niche breadth and aridity

After evaluating multiple models/transformations in the phylogenetic linear regression analyses, the Lambda transformation consistently provided the best fit for examining the covariation between niche breadth (defined as the central 95% variation along the first two principal components) and species positions along the aridity gradient (median values; Supplementary Table 1). Consequently, the interpretations presented below are based on the results using Lambda.

In line with the notion of climate specialization, across all regression models and aridity proxies, the relationship between climatic niche breadth and aridity is consistently negative (Figure 2; Supplementary Figure 10; Supplementary Table 2), even when controlling for log-range area (Supplementary Figure 11; Supplementary Table 3; Supplementary Table 4) and with the alternative aridity metric (percentage of arid cells in species’ distribution ranges; Supplementary Figure 12, Supplementary Table 5). However, we observe variation in the steepness of the slopes among clades (Supplementary Tables 2-5). Across all proxies of aridity, for both niche breadth of PC1 and PC2, squamates and birds tend to show the steepest slopes in every analysis, indicating the strongest climatic specialization under arid conditions, while slopes tend to be shallower for amphibians and mammals.

**Figure 2.**
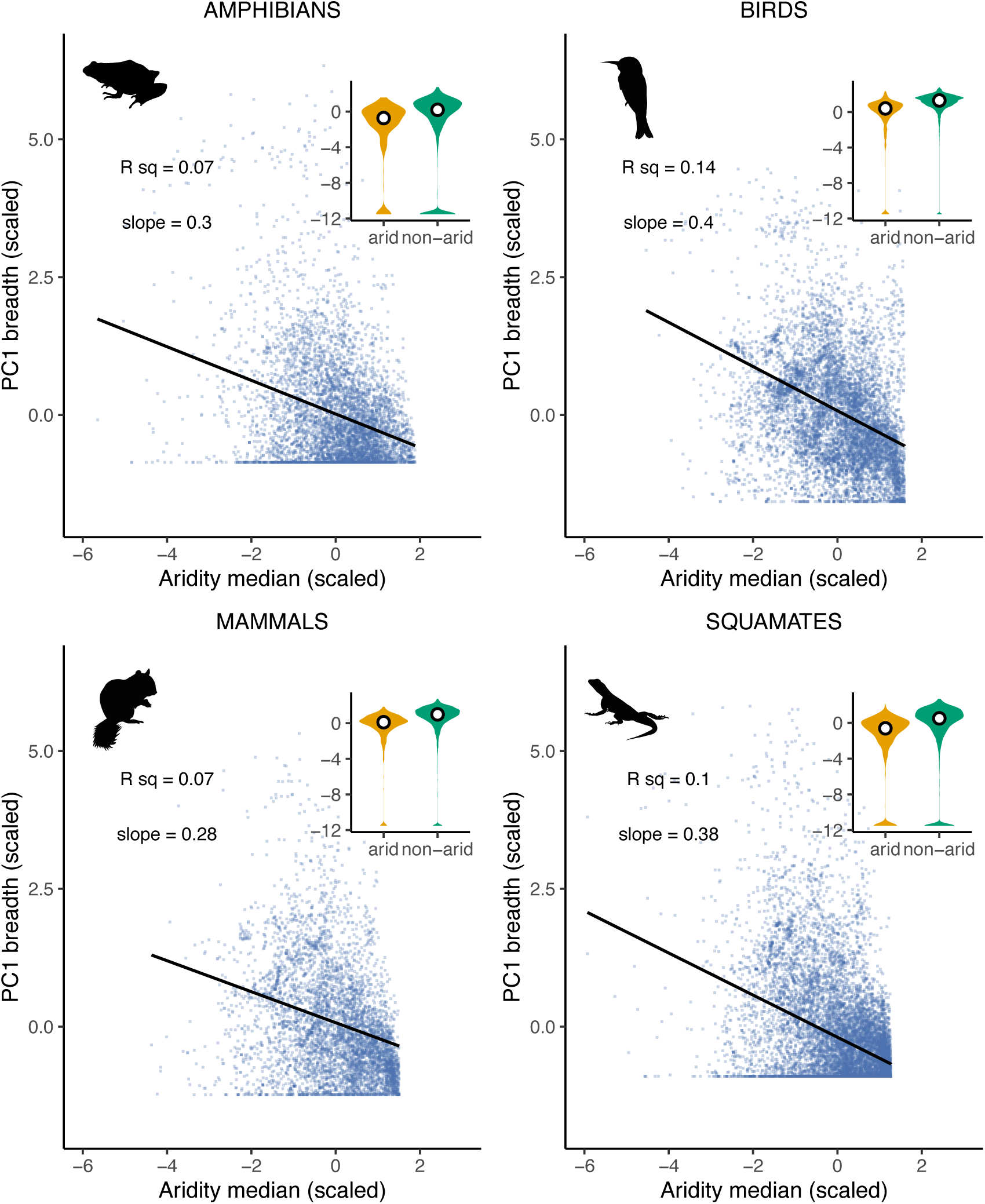
Relationship between species’ median aridity and climatic niche breadth (PC1). There is a pervasive negative relationship (i.e., arid species have generally narrower niche breadth than non-arid species), although the strength of that relationship varies across tetrapod groups.

This negative relationship between niche breadth and aridity was also detected when studied for each desert independently (Supplementary Figure 13). Interestingly, we found that the slopes of the within-pool regression models were very similar in the two endotherms groups (birds and mammals), differing substantially from those of amphibians and squamates, which in turn showed important similarities in some deserts but notable differences in others (Supplementary Figure 14). Likewise, for the four groups of tetrapods, we found significantly lower niche breadths in arid species relative to non-arid species in the phylogenetic ANOVA analyses (Supplementary Figure 15; Supplementary Table 6).

### Analyses of Phylogenetic Clustering

#### Large-scale analysis

Consistently with the idea of increased habitat filtering in arid environments, with the exception of birds, we detected significantly lower levels of Faith’s phylogenetic diversity (PD) in the deserts than in the simulated assemblages including the surrounding non-arid regions (Figure 3).

**Figure 3.**
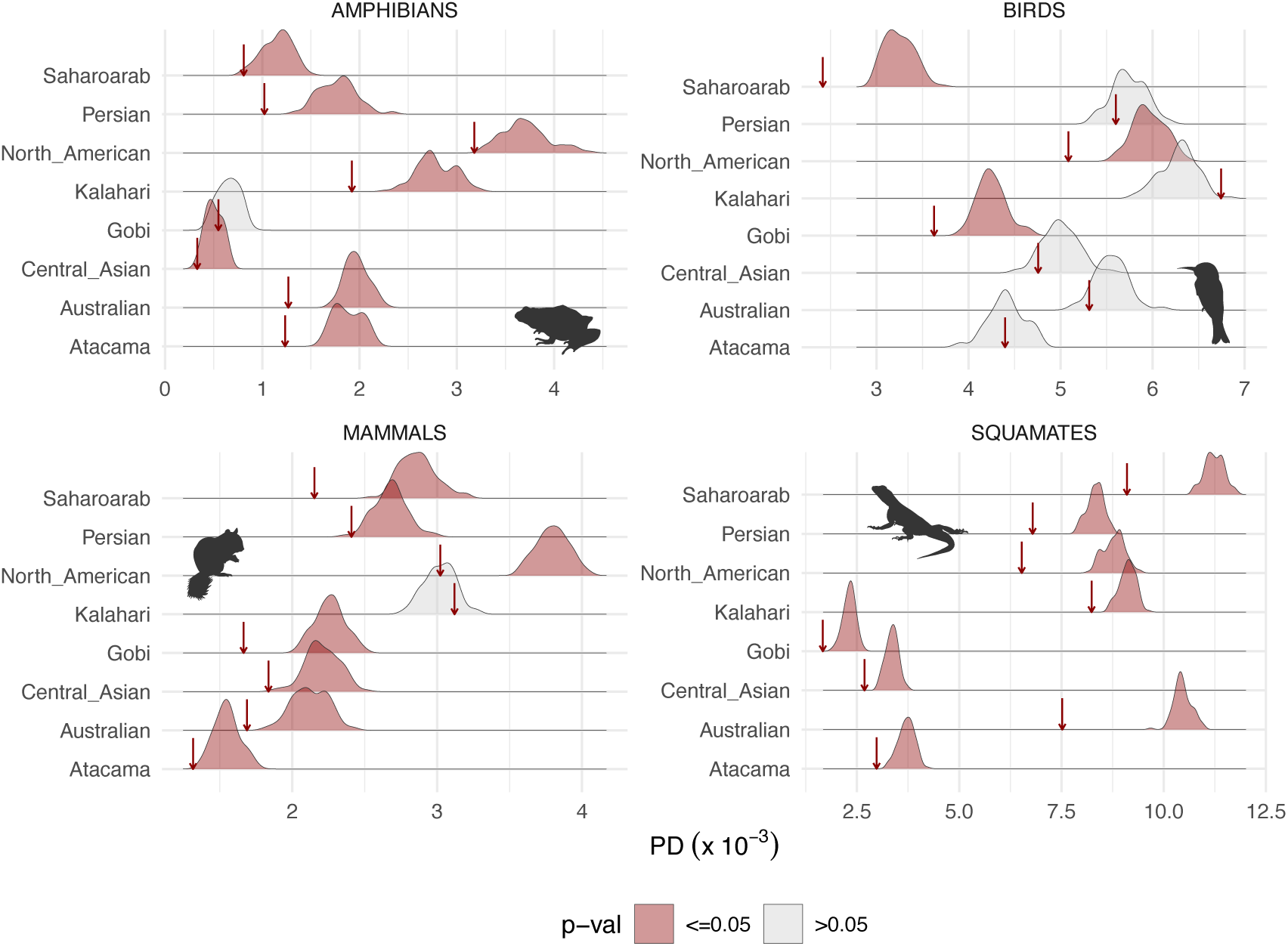
Observed Faith’s phylogenetic diversity (PD) in the arid assemblages (red arrows) compared to a null model based on the PD of 100 species pools constituted by random permutations of species present in the buJer regions surrounding each arid system (see Supplementary Figure 2).

We identified almost identical results with mean nearest taxon distance (MNTD; Supplementary Figure 16), with significantly lower than expected levels in almost every desert assemblage (except for birds) indicating that there are very closely related species in the arid communities. On the contrary, we found that the mean pairwise distance (MPD) is not lower than expected in many desert assemblages, not only in birds (as in the PD and MNTD analyses), but also especially in squamates (Supplementary Figure 17), reflecting that not all species in these desert communities are very closely related. These results suggest that desert communities of amphibians, squamates and mammals are indeed composed of phylogenetically clustered species overall (low PD and MNTD levels), but these closely related species belong to phylogenetically dispersed clades (non-significant MPD levels). In other words, consistent with the idea that an ecological filter exists, only a few clades persist in the desert, though they can be in relatively distant parts of the phylogeny.

#### Regional (cell-based) distribution of phylogenetic diversity

Residual PD is not uniformly distributed across the world, including arid regions, and the patterns differ among tetrapod groups (Figure 4; Supplementary Figure 18; Tejero-Cicuéndez et al. 2025). Indeed, our results show that different arid regions exhibit different and even opposing PD diversity patterns, and that these patterns are clade-specific. For instance, Australia generally has high residual PD for birds and mammals, but low residual PD for amphibians and squamates. The South African arid region, encompassing the Kalahari desert, shows overall high levels of residual PD across tetrapods, except for some areas with low amphibian residual PD. The Sahara desert harbors high residual PD for squamates but contains areas of both high and low residual PD for mammals. For birds, the Sahara does not show particularly low or high residual PD, but the sub-Saharan belt stands out with the highest bird residual PD. For amphibians, which do not thrive in the Sahara Desert, the sub-Saharan belt is instead a region of low residual PD.

**Figure 4.**
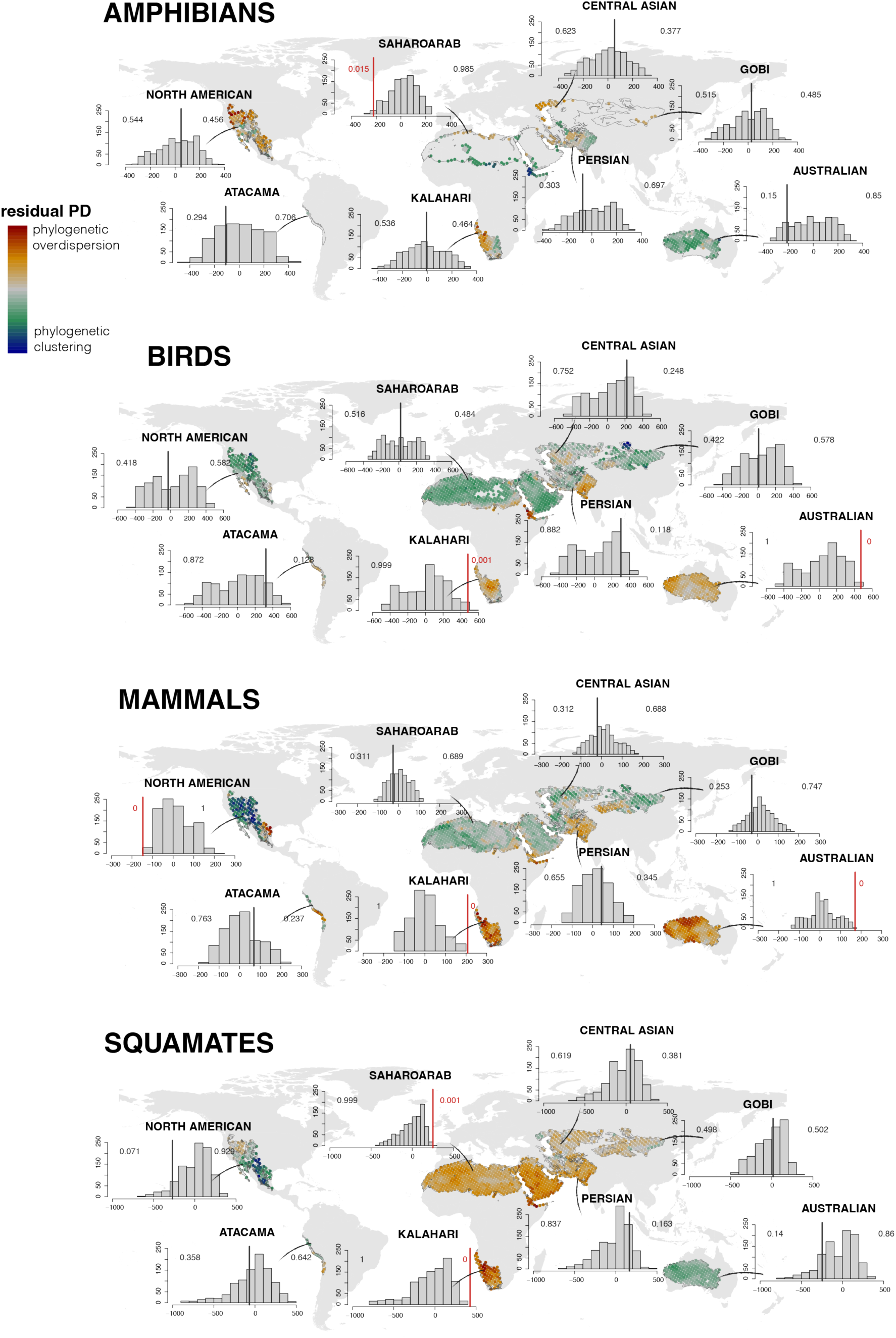
Geographic patterns of residual phylogenetic diversity (residual PD) in the deserts for each tetrapod group. In each desert, the comparison between the observed average residual PD (vertical lines) and the obtained through the null model (gray bars) is shown.

In birds and squamates, the observed average residual PD in the 25% most arid locations is significantly higher (right p-value: birds = 0.01, squamates = 0.035) than the obtained from the null hypothesis (999 rotations) (Supplementary Figure 19). In amphibians, we found significantly low levels of residual PD in the 25% most arid locations (left p-value = 0.039). In mammals, we did not find a particularly high or low residual PD in the most arid locations relative to the null hypothesis.

Per-desert regional PD inspection provided further insight into these finer scale results. In amphibians, the overall low levels of residual PD are driven by the species assemblages from the Saharo-Arabian desert, while there are no significantly high or low residual PD values in the rest of the arid systems (Figure 4). In birds, high overall residual PD results from regional communities in the Kalahari and the Australian deserts, while in squamates they reflect communities in the Kalahari and the Saharo-Arabian deserts. Lastly, the non-significant overall results for mammals actually hide significantly higher than expected residual PD in the Australian and Kalahari deserts, and lower than expected in the North American arid-adapted communities. This reflects the heterogeneity among desert systems, and suggests that phylogenetic overdispersion (higher residual PD than expected) at relatively small scales is an important pattern in several arid-adapted vertebrate communities.

## Discussion

This study provides the first global perspective on how drylands act as drivers of ecological specialization and ecological filters for terrestrial vertebrates. Consistent with the notion that arid regions impose strong constraints on biodiversity, our findings confirm that deserts harbor relatively low vertebrate diversity overall, despite covering nearly 40% of Earth’s land surface (Gaur and Squires 2018). Yet this pattern is not uniform across clades: while in amphibians less than 4% of their diversity is associated to arid regions, other groups, most notably squamates, are frequently associated to arid conditions, with more than 20% of their global diversity found in drylands. This striking disparity underscores that deserts are both strong ecological filters and important areas of colonization and diversification, depending on lineage.

Despite these differences in absolute diversity, all four tetrapod groups exhibited a pervasive negative relationship between climatic niche breadth and aridity, even after accounting for range size (Figure 2; Supp. Figures 10-12, 15). This supports the idea that deserts demand specialized and evolutionarily costly adaptations, which in turn constrain species to arid environments and limit their ecological flexibility (Williams and Tieleman 2002; Cox and Cox 2015). This is consistent with previous studies that found that climatic niche breadth is narrower in terrestrial vertebrates inhabiting warmer and drier regions (Bonetti and Wiens 2014; Lin and Wiens 2017; Lin et al. 2019; Pie et al. 2021).

The intensity of this specialization, however, varies among clades (Figure 2). In general, birds and squamates show steeper relationships between aridity and niche breadth, whereas mammals and amphibians show shallower slopes. These differences likely reflect both physiological constraints and the repertoire of behavioral strategies available to each group. For instance, mammals may buffer environmental extremes through behaviors such as nocturnality, burrowing, or physiological mechanisms like adaptive heterothermy and the production of concentrated urine and dry feces (Nagy 2004; Costa et al. 2013; Fuller et al. 2016), strategies that are less frequent in birds and may reduce the need for extreme specialization. In amphibians, a combination of strategies such as nocturnality, burrowing, and niche tracking, often restricted to rare wet habitats within arid landscapes (e.g., oases), may explain their comparatively shallower slope (McClanahan et al. 1994; Davis et al. 2013; Zylstra et al. 2015). By contrast, squamates include a high diversity of heliothermic species that are tightly linked to external thermal environments (Garcia-Porta et al. 2019; Dubiner et al. 2024), increasing the need for physiological and morphological adjustments in arid regions. Moreover, the presence of massive adaptive radiations in some deserts, such as the Australian system (Blom et al. 2016), may further amplify the observed patterns of niche specialization in this group.

These clade-level patterns also interact with geography. Endotherms (birds and mammals) tend to show relatively consistent responses across desert systems, whereas ectotherms (squamates and amphibians) vary more strongly among regions (Supp. Figure 14). This likely reflects both organismal traits, such as ectotherms’ direct dependence on external thermal regimes, and the ecological and evolutionary idiosyncrasies of individual deserts, which differ in resource availability, microclimatic refugia, substrate types, and evolutionary age (Tejero-Cicuéndez et al. 2022).

At the community level, our analyses revealed that most arid faunas exhibit lower phylogenetic diversity and stronger clustering than expected under null models (Figure 3), consistent with ecological filtering and the stress-dominance hypothesis (Ramm et al. 2018). This was evident in Faith’s PD and MNTD (Supplementary Figure 16), while MPD values were often not significantly different from random expectations (Supp. Figure 17). This combination suggests that deserts are typically composed of clusters of closely related species, but that these clusters may arise from multiple, independent colonizations by distinct clades, rather than a single evolutionary radiation. Squamates exemplify this pattern, with many independent (and distantly related) radiations contributing to their diversity in deserts (e.g., Rabosky et al. 2007; Šmíd et al. 2013; Wiens et al. 2013; Blair and Sánchez-Ramírez 2016; Blom et al. 2016; Tamar et al. 2016; Barley et al. 2019; Simó-Riudalbas et al. 2019; Jennings 2021).

Birds stand out as an exception to this general pattern. Despite their pronounced climatic specialization, they do not exhibit strong phylogenetic clustering at large scales (Figure 3). We interpret this as a consequence of their high dispersal capacity. This likely facilitates repeated, independent colonization of arid systems and long-distance movements among deserts, resulting in a mixing of lineages that decreases phylogenetic clustering at the system level (Claramunt et al. 2012; Weeks and Claramunt 2014; Saito et al. 2015; Suárez et al. 2022). By contrast, mammals, amphibians, and squamates show stronger clustering, consistent with more clade- and region-specific colonization that enhances phylogenetic structuring.

Importantly, phylogenetic patterns proved scale-dependent. At large scales, deserts act as ecological filters, producing clustered assemblages, while at regional scales, clustering often weakens or reverses into phylogenetic overdispersion (Figure 4). This reflects a shift in assembly processes: environmental constraints dominate broad-scale patterns, whereas at finer scales, other processes such as competition, dispersal, and microhabitat partitioning become more influential (Webb et al. 2002; Cavender-Bares et al. 2009). These results echo the broader view that “environmental filtering” is not a single, uniform mechanism but the outcome of interacting abiotic and biotic processes (Kraft et al. 2015; Cadotte and Tucker 2017).

Nonetheless, exceptions highlight the heterogeneity of desert systems. Amphibians in Saharo-Arabia and mammals in North America, for instance, retain significantly low PD even at regional scales, likely reflecting either exceptionally strong environmental constraints or the imprint of local adaptive radiations.

Taken together, our results demonstrate that drylands function as powerful drivers of climatic specialization and ecological filtering, but that their imprint on biodiversity is clade- and scale-dependent. Birds and squamates show particularly strong specialization to arid conditions, while mammals and amphibians appear more buffered. Future mechanistic studies linking phylogenetic patterns to functional traits and demographic performance will be essential to disentangle abiotic constraints from biotic interactions and to uncover the adaptations that enable persistence in deserts (Lara-Reséndiz et al. 2022). Finally, our findings carry urgent conservation implications. First, arid species with narrow niche breadths may face a heightened risk of extinction (Grinder and Wiens 2023). This is particularly concerning because arid regions have historically received less conservation attention than hyperdiverse tropical systems, despite their unique and highly specialized biodiversity (Brito and Pleguezuelos 2020). Second, as drylands continue to expand and climatic extremes intensify (Harris et al. 2018), the pace of environmental change may exceed the ability of mesic-adapted species to evolve the necessary specializations, constraining their capacity to persist in these increasingly widespread habitats. Therefore, understanding how the expansion of arid regions, and the human impacts within them, amplifies biodiversity vulnerability both within and beyond deserts is crucial for guiding actions that mitigate the ongoing global extinction crisis.

## Data and code availability statement

All data used for the analyses presented here is published elsewhere and the original sources indicated in the text. All the scripts developed and used for the analyses and figures are available in a Zenodo repository, publicly available upon publication.

## Disclosure on the use of Artificial Intelligence

We used a generative large language model (OpenAI ChatGPT) to assist with wording refinements in specific parts of the manuscript and with R code drafting; all outputs were reviewed, verified, and edited by the authors. All scientific content, study design, analyses, and interpretations were conceived and performed solely by the authors.

## Acknowledgments and Funding

We are grateful to Dean C. Adams and Juan G. Rubalcaba for their help with statistical analysis. HT-C was supported by a “Juan de la Cierva - Formación” postdoctoral fellowship (FJC2021-046832-I) from the Spanish MCIN/AEI/10.13039/501100011033 and the European Union NextGenerationEU/PRTR, and by a Humboldt Postdoctoral Fellowship from the Alexander von Humboldt Foundation. IM was funded by the Alexander von Humboldt Foundation through a Humboldt Postdoctoral Fellowship. GA was funded by the US National Science Foundation award number 2020424: “AccelNet: International Tropical Forest Science Alliance (ITFSA): A multi-network science and training initiative to accelerate understanding of the role of tropical forests in the Earth System”. JG-P was supported by the program “Atracción de Talento Investigador Modalidad I” from the Spanish Comunidad de Madrid (2022-T1/AMB-24171).

## Notes

### Competing Interest Statement

The authors have declared no competing interest.

