## Supplementary Figure for "Aridity Drives Climatic Specialization and Phylogenetic Clustering in Terrestrial Vertebrates"

a)

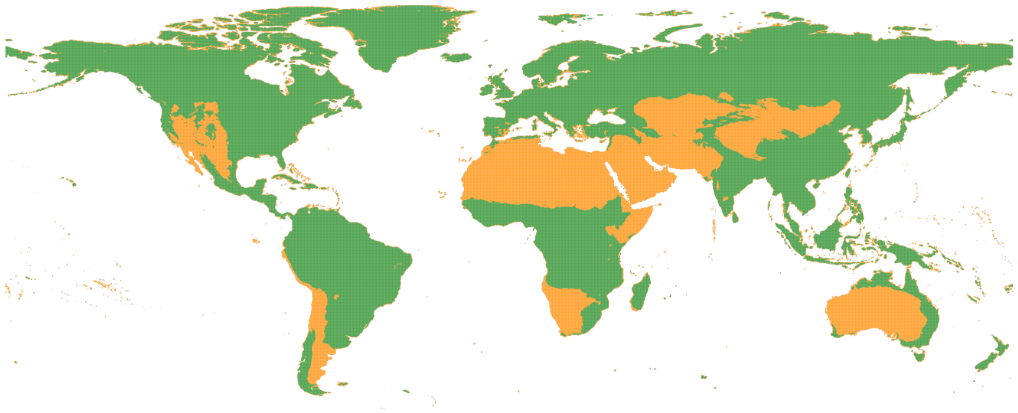

b)

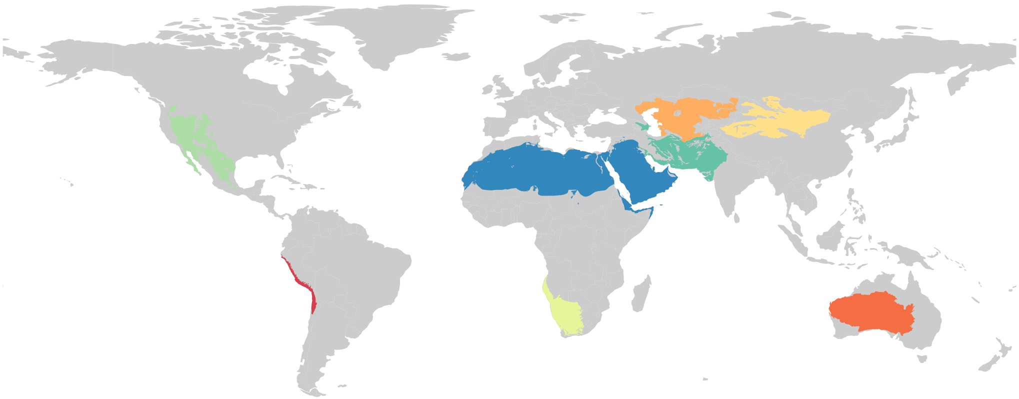

**Supplementary Figure 1**. Comparison of the two approaches used to define aridity and categorize arid species. a) Map of arid regions based on the Aridity Index (arid regions are here defined as AI < 0.25). b) Map of the major arid systems in the world, based on the Biome 13 in Olson et al. (2001): Desert and xeric shrublands.

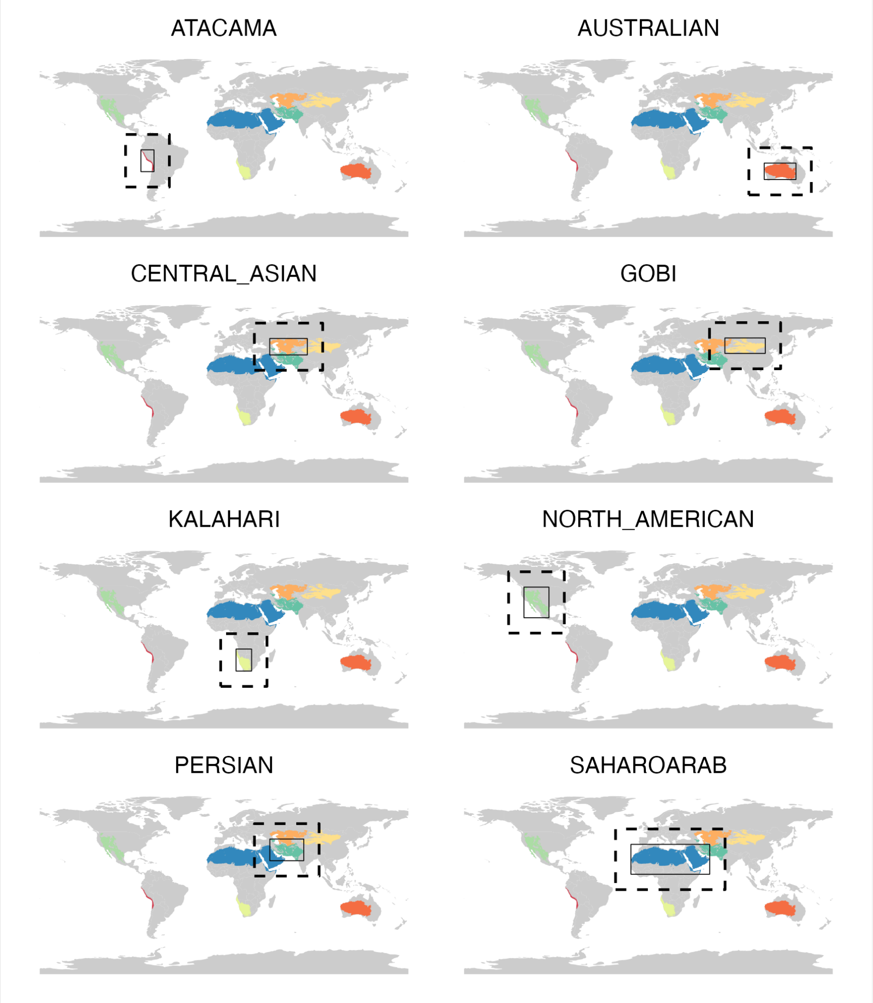


**Supplementary Figure 2**. Buffers surrounding each arid system, used for building species pools and comparing arid-adapted communities to non-arid assemblages.

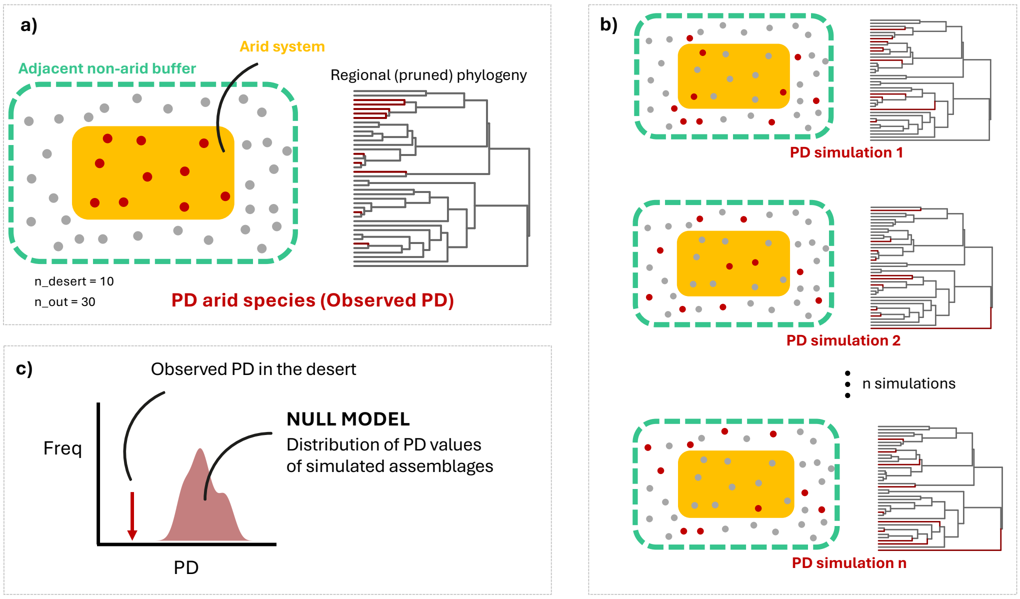


**Supplementary Figure 3**. Conceptual visualization of the large-scale phylogenetic diversity (PD) analyses by generating a null model to assess the significance of PD values in each arid system, comparing the observed value of the communities within the deserts (a) to n simulated assemblages of species within the buffers surrounding each desert (b) (See above Supplementary Figure 2). The comparison shows the significance of the observed values relative to the null model (c).

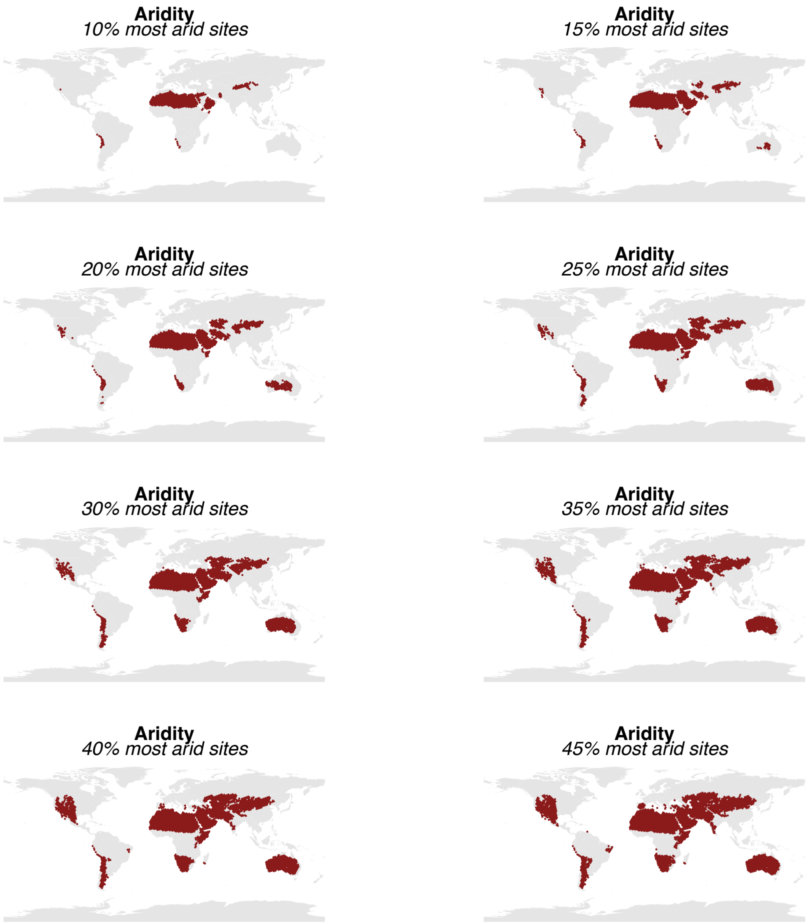


**Supplementary Figure 4**. Map of the 270x270km locations with highest aridity, with different thresholds (from 10% to 45%).

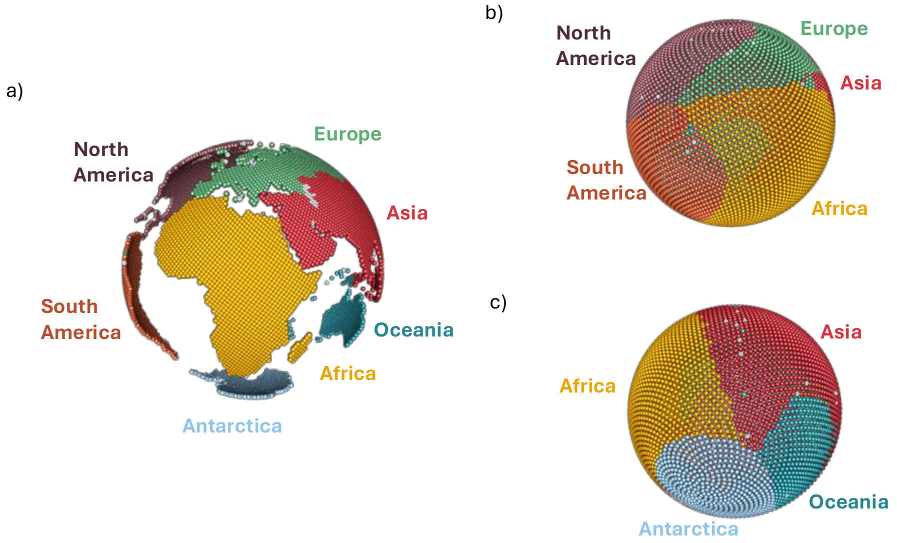


**Supplementary Figure 5**. One-to-one assignment of uniformly distributed locations within the continents (a) and uniformly distributed locations completely covering a hypothetical “only-land” Earth (b and c). b) and c) show two different angles, to facilitate the understanding of this step in our workflow. This step allows us to rotate geographically distributed variables (e.g., aridity index), avoiding the subsequent problem of the oceans resulting from studying terrestrial diversity patterns.

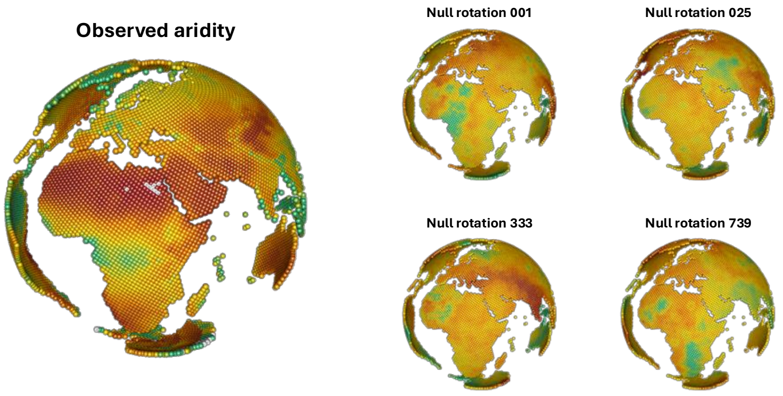


**Supplementary Figure 6**. Visualization of the observed geographic patterns of aridity, and the null patterns resulting from four of the 999 random 3D rotations implemented to build the null model (see main text for further details). In each rotation, the geographic distribution of the environmental variable changes, and we built the null model by registering the phylogenetic diversity (residual PD) in the most arid sites after each rotation, and then comparing it to the observed value of residual PD in the real most arid sites.

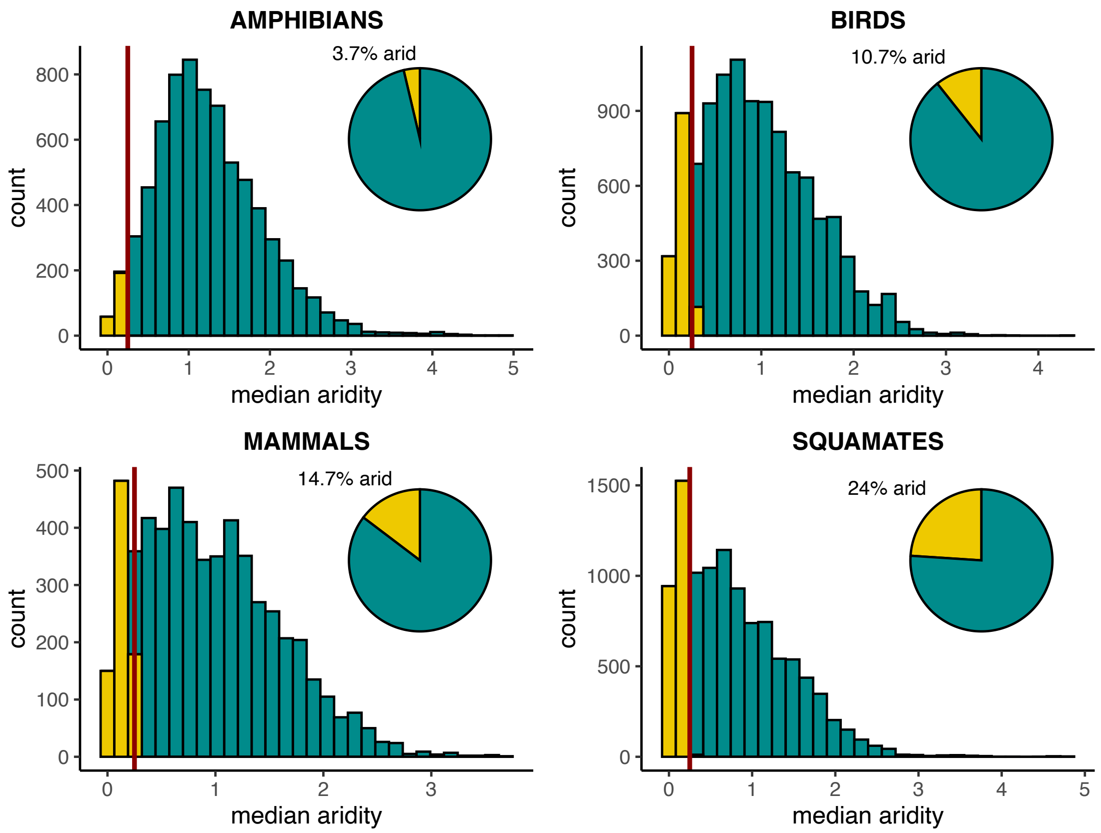


**Supplementary Figure 7**. Percentage of arid species in each tetrapod clade, based on the median value of the aridity index (AI) in species' distribution ranges. A species was considered 'arid' when its median AI < 0.25 (see map in Supplementary Figure 1a).

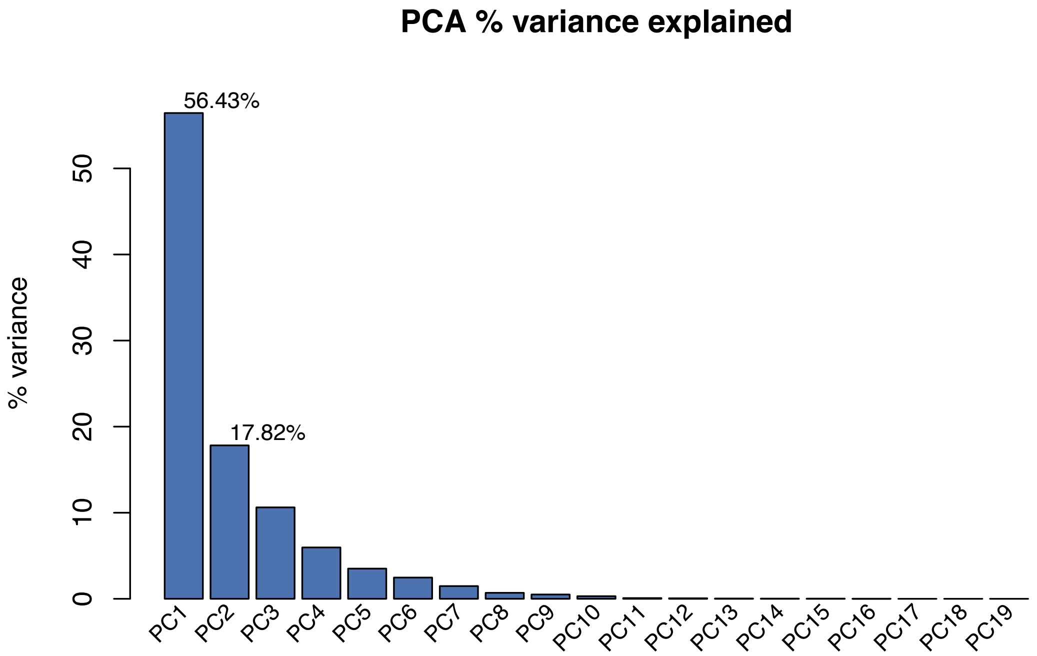


**Supplementary Figure 8**. Variance explained by the different principal components in the environmental PCA.

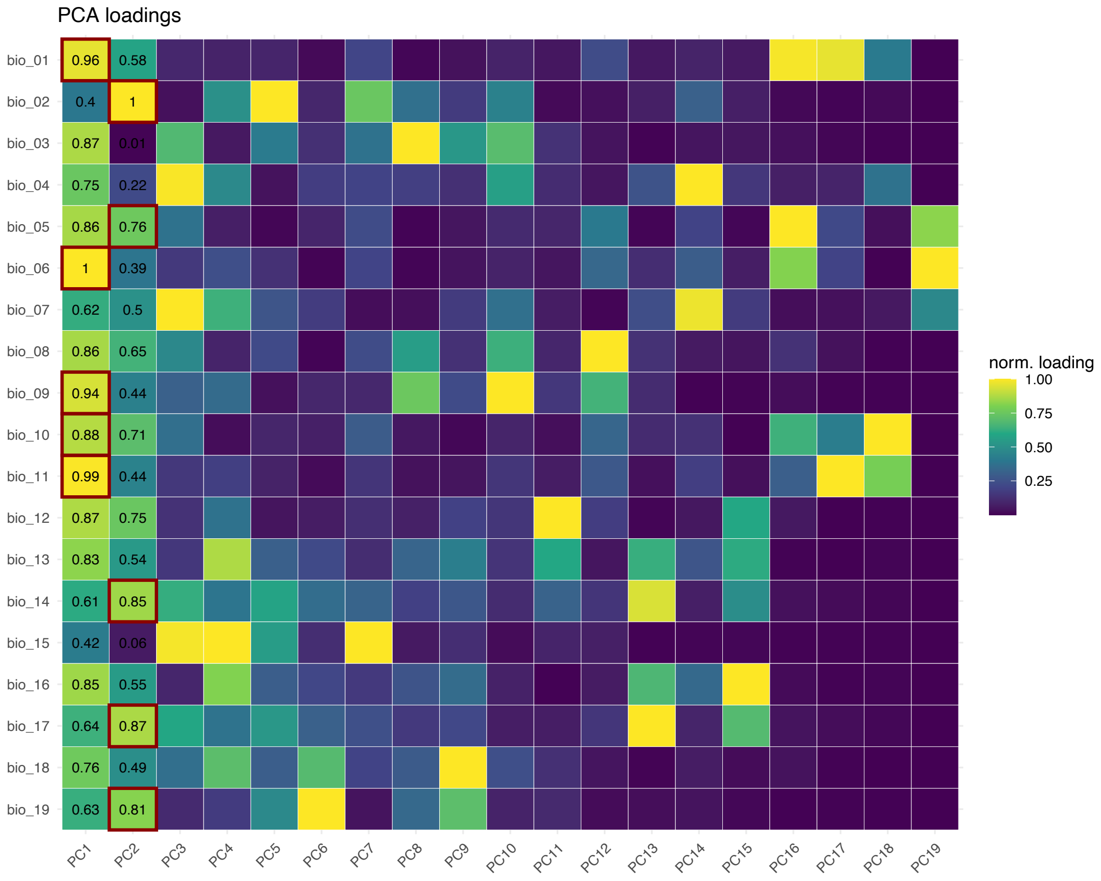


**Supplementary Figure 9**. Loadings of the Principal Component Analysis (PCA) showing the most important variables in each principal component.

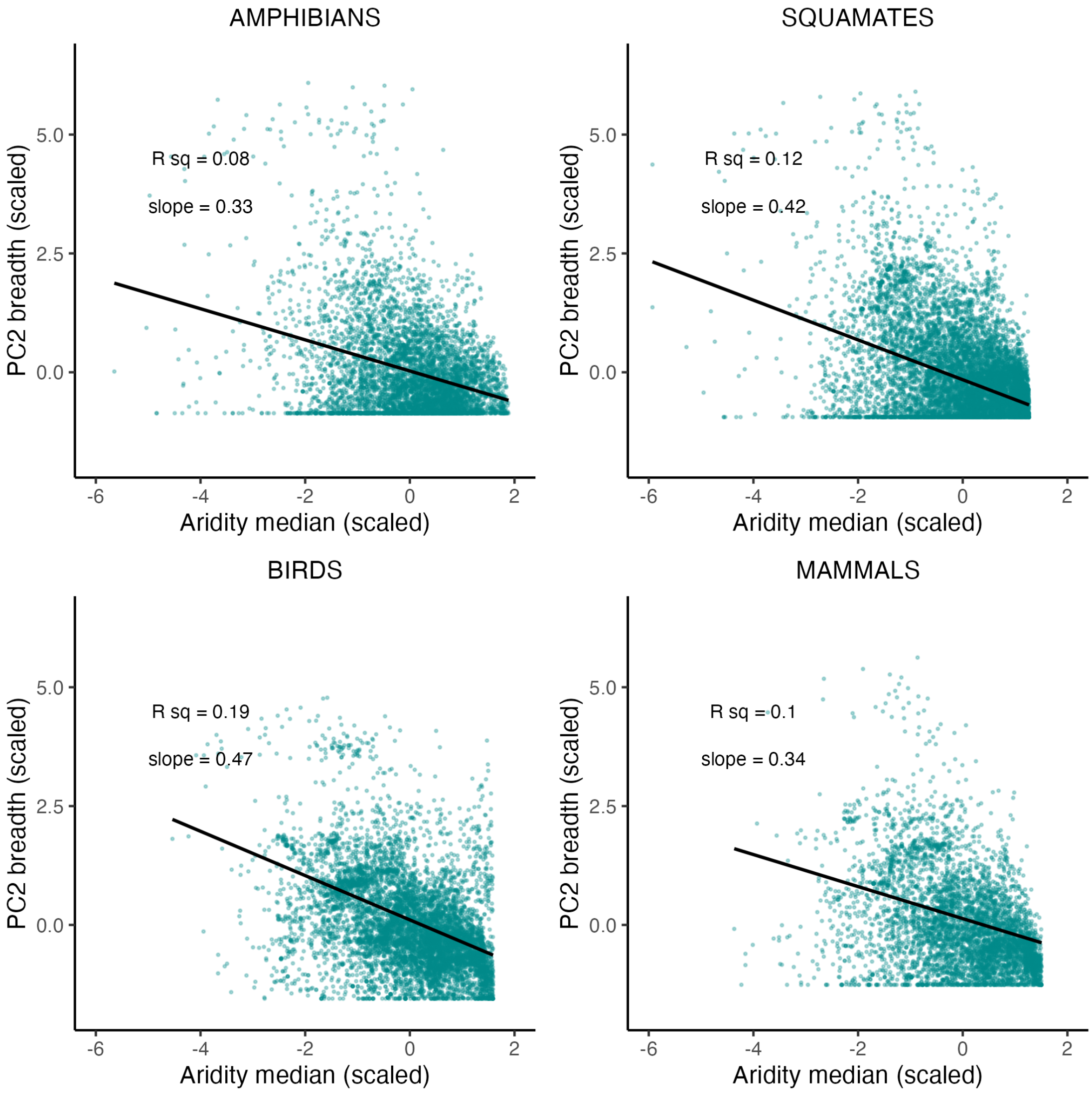


**Supplementary Figure 10**. Relationship between species' median aridity and climatic niche breadth (PC2).

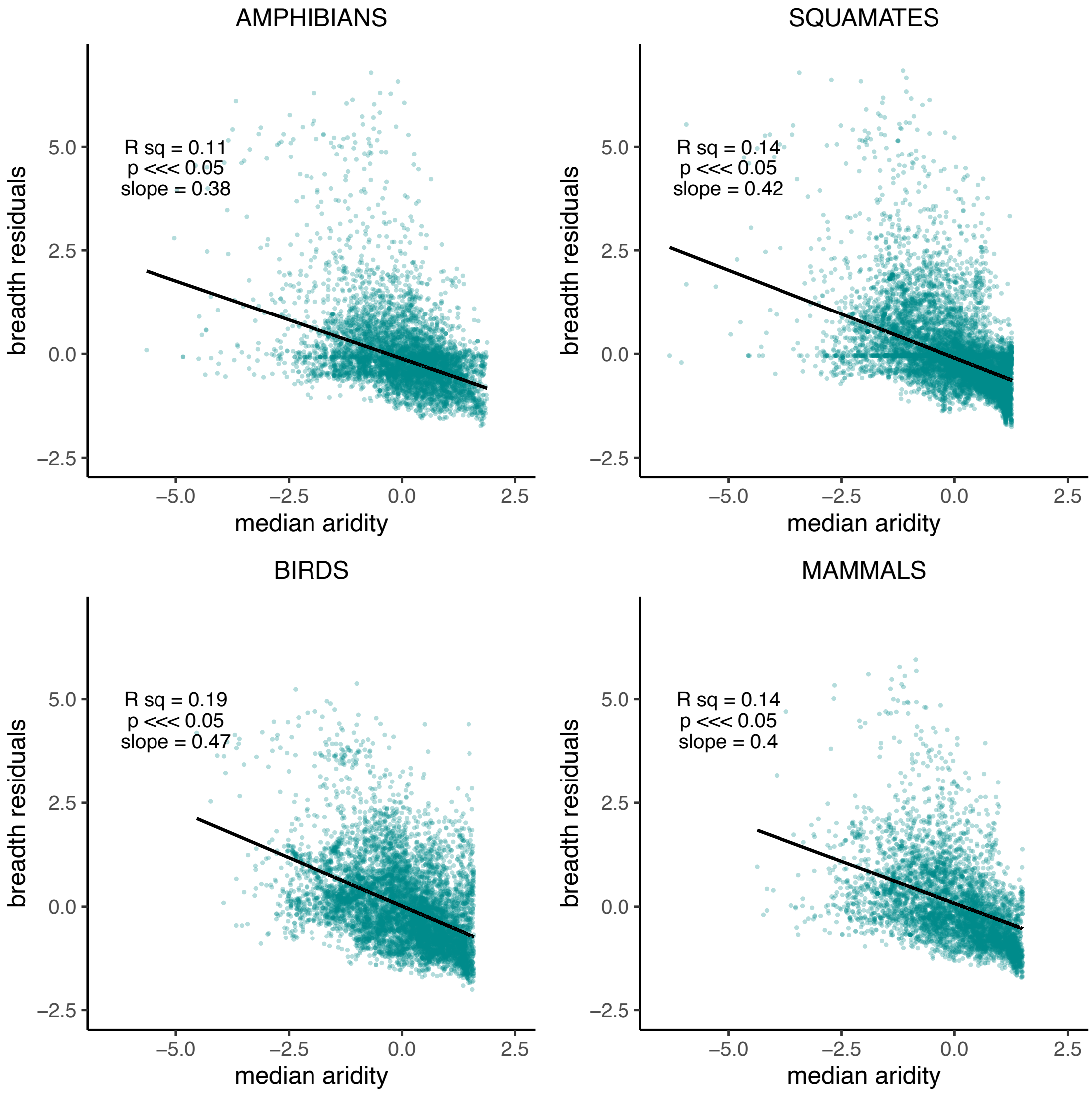


**Supplementary Figure 11**. Regression between species' median aridity and niche breadth (PC1) after correcting for log-range area.

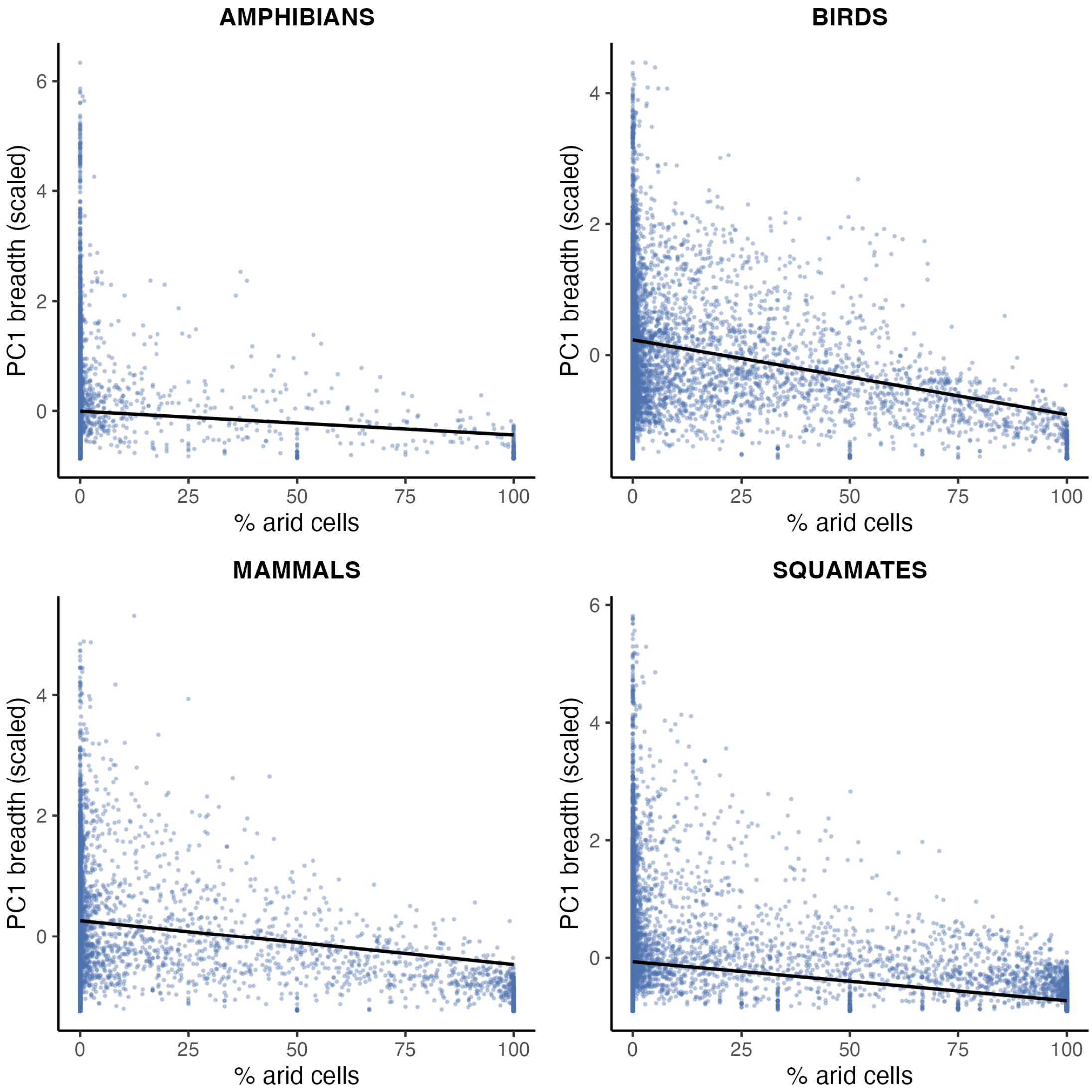


**Supplementary Figure 12**. Regression between the percentage of arid cells in species' distribution ranges and climatic niche breadth (PC1).

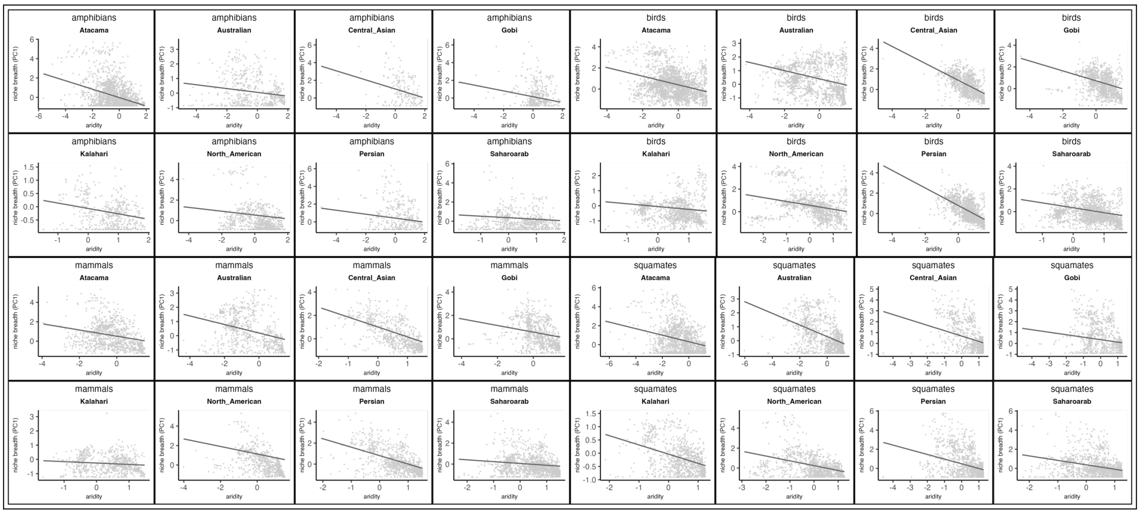


**Supplementary Figure 13**. Relationship between median aridity and climatic niche breadth (PC1) for each arid system independently.

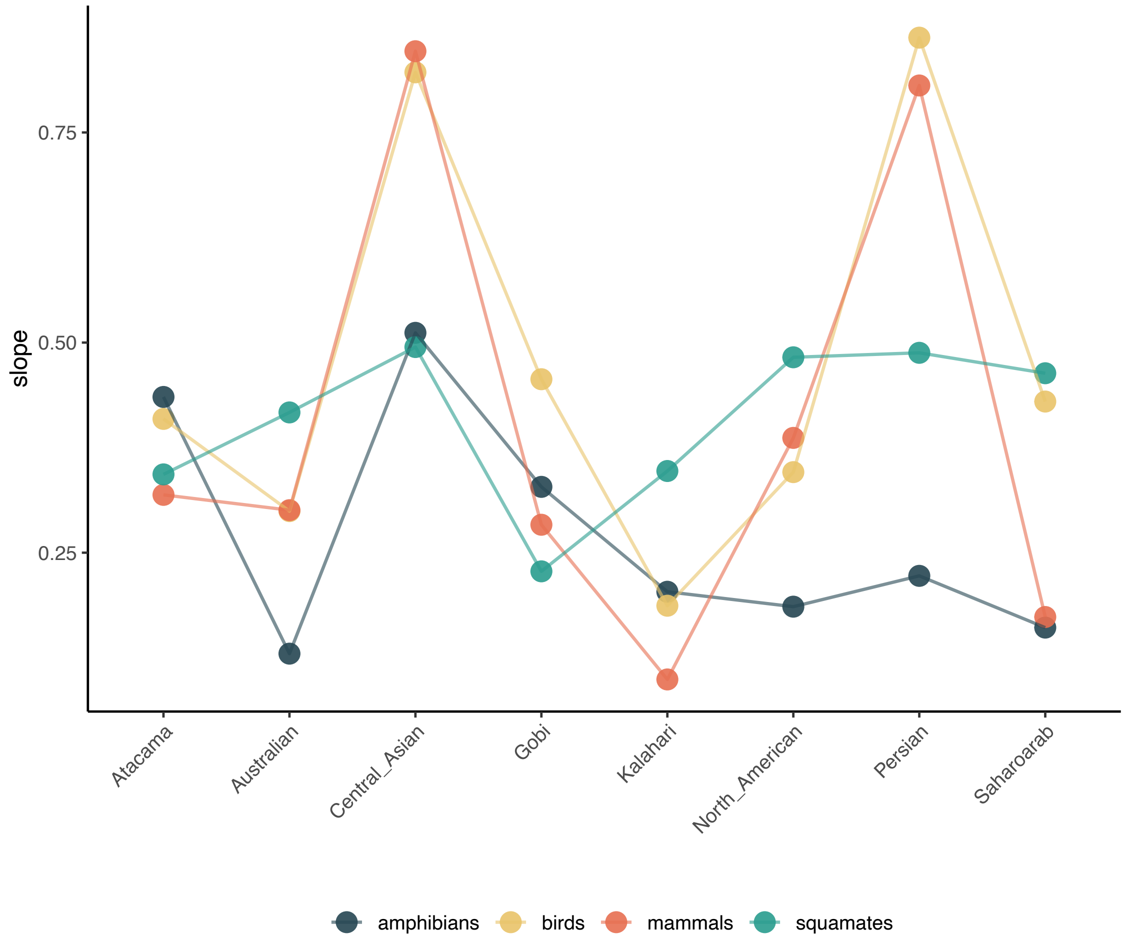


**Supplementary Figure 14**. Absolute slope values from the regression models testing the relationship between median aridity and climatic niche breadth (PC1) for each arid system independently. There is a conspicuous pattern in the two endotherms groups (birds and mammals), while ectotherms (amphibians and squamates) are similar in some deserts but very different in others.

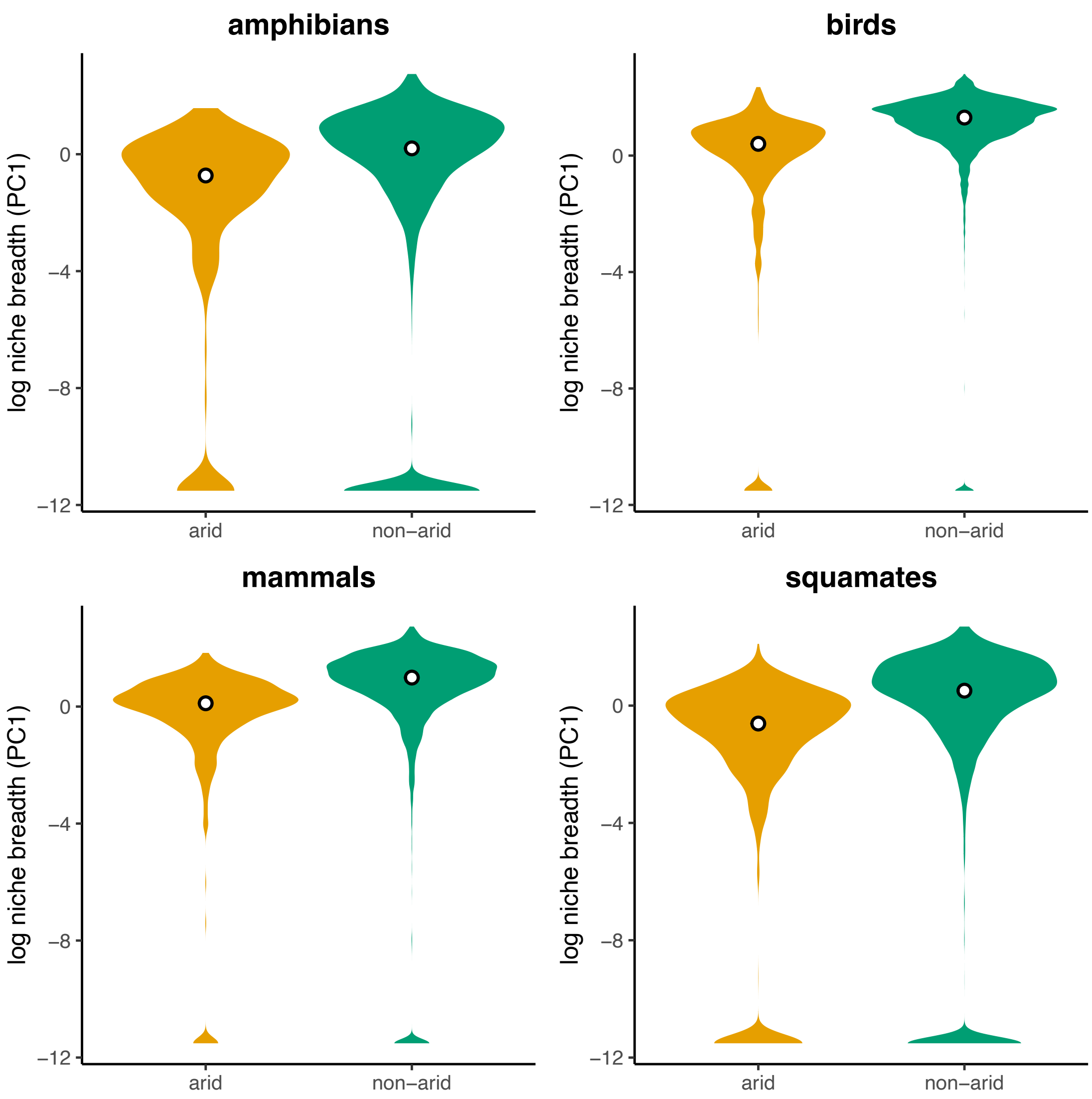


**Supplementary Figure 15**. Differences in log-transformed climatic niche breadth (PC1) between arid and non-arid species.

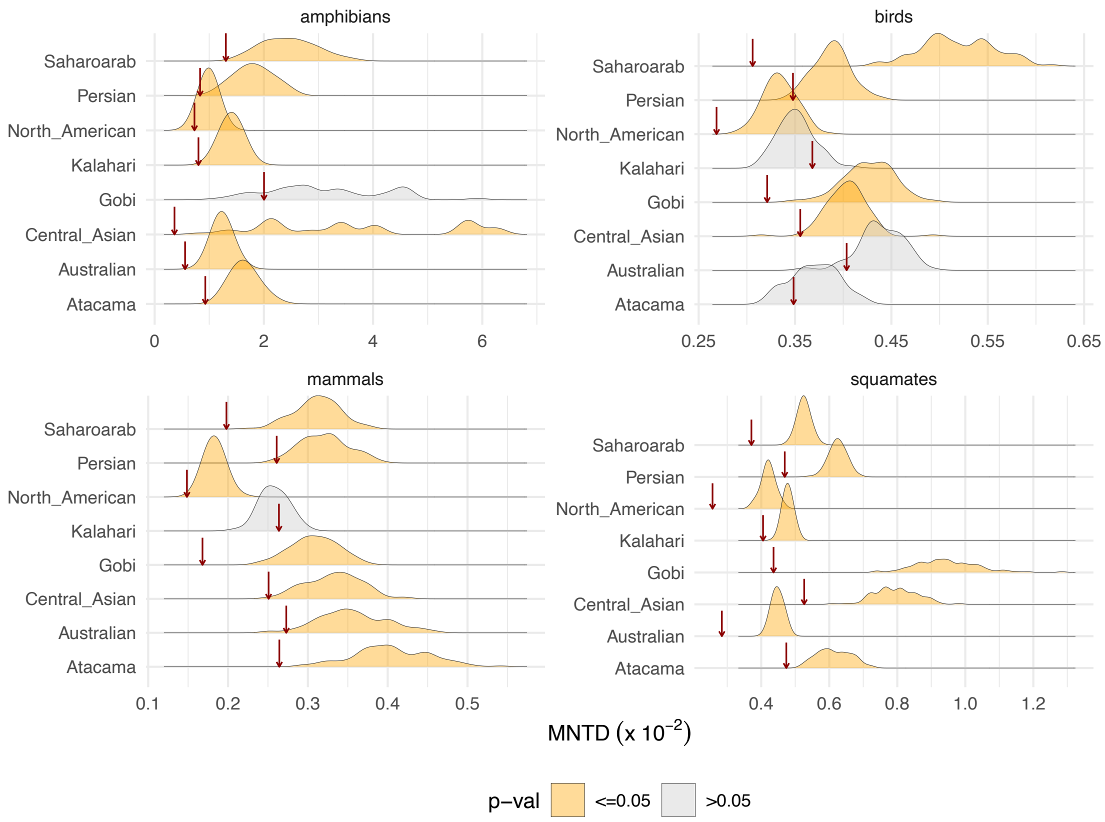


**Supplementary Figure 16**. Mean nearest taxon distance (MNTD) in arid-adapted communities (red arrows) compared to a null model based on the MNTD of 100 species pools constituted by random permutations of species present in the buffer regions surrounding each arid system (see Supplementary Figure 2).

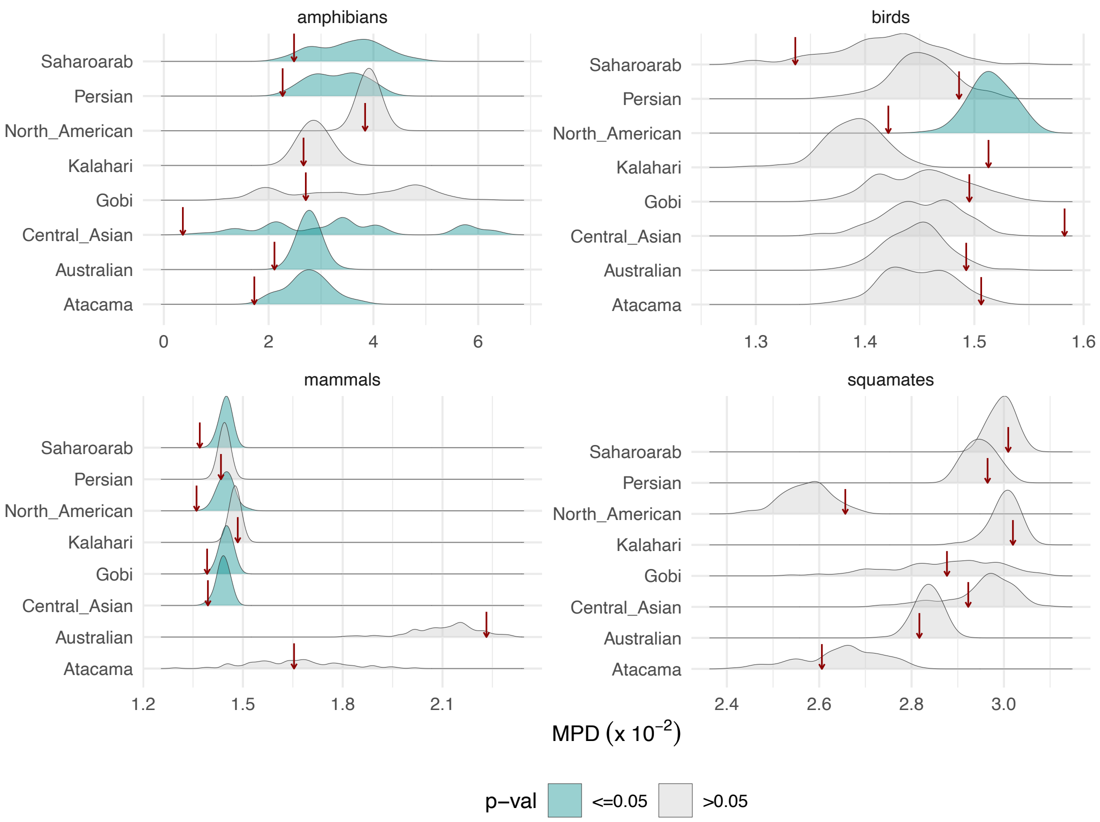


**Supplementary Figure 17**. Mean pairwise distance (MPD) in arid-adapted communities (red arrows) compared to a null model based on the MPD of 100 species pools constituted by random permutations of species present in the buffer regions surrounding each arid system (see Supplementary Figure 2).

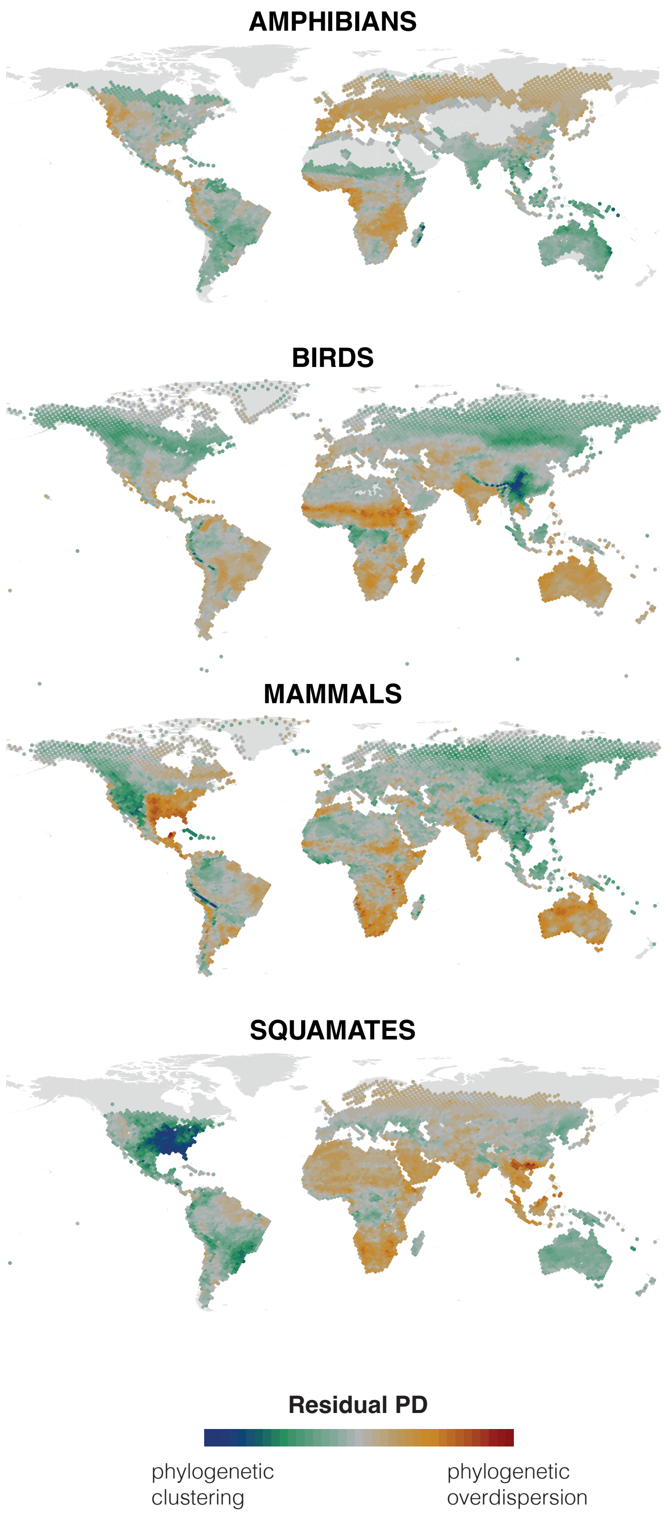


**Supplementary Figure 18**. Global geographic patterns of residual phylogenetic diversity (residual PD) in the four groups of tetrapods.

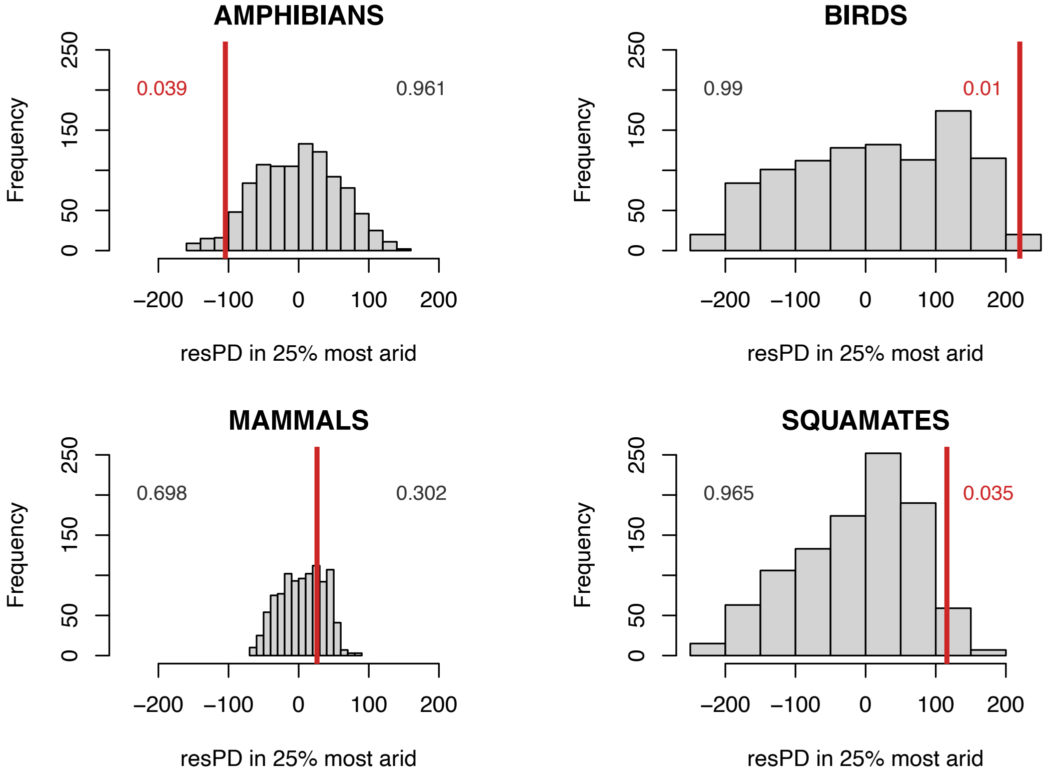


**Supplementary Figure 19**. Observed (red lines) and simulated (gray bars) average residual PD values in the 25% most arid locations for each tetrapod group. Numbers in red indicate significant differences.
