## Supplementary Table for "Aridity Drives Climatic Specialization and Phylogenetic Clustering in Terrestrial Vertebrates"

**Supplementary Table 1**. Model selection among the evolutionary models and transformations tested for the phylogenetic regression between niche breadth (PC1 and PC2) and median aridity. Lambda is the best supported in all cases. OUr: Ornstein-Uhlenbeck random root; OUf: Ornstein-Uhlenbeck fixed root; BM: Brownian Motion; EB: Early Burst.

| | | PC1 breadth | | | PC2 breadth | | |
|---|---|---|---|---|---|---|---|
| | model | fit | delta | w | fit | delta | w |
| amphibians | lambda | 15589.66 | 0 | 1 | 15516.27 | 0 | 1 |
| | OUr | 16028.26 | 438.5974 | 5.75E-96 | 15969.59 | 453.3261 | 3.6E-99 |
| | OUf | 16028.26 | 438.5974 | 5.75E-96 | 15969.59 | 453.3261 | 3.6E-99 |
| | BM | 18819.09 | 3229.425 | 0 | 18766.84 | 3250.57 | 0 |
| | EB | 18821.09 | 3231.425 | 0 | 18768.84 | 3252.57 | 0 |
| squamates | lambda | 23852.44 | 0 | 1 | 23835.01 | 0 | 1 |
| | OUr | 24647.2 | 794.766 | 2.6E-173 | 24634.43 | 799.4205 | 2.6E-174 |
| | OUf | 24647.2 | 794.766 | 2.6E-173 | 24634.43 | 799.4205 | 2.6E-174 |
| | BM | 29414.45 | 5562.015 | 0 | 29929.72 | 6094.709 | 0 |
| | EB | 29416.45 | 5564.015 | 0 | 29931.72 | 6096.709 | 0 |
| birds | lambda | 20604.41 | 0 | 1 | 20012.76 | 0 | 1 |
| | OUr | 21258.3 | 653.8913 | 1E-142 | 20732.78 | 720.014 | 4.5E-157 |
| | OUf | 21258.3 | 653.8913 | 1E-142 | 20732.78 | 720.014 | 4.5E-157 |
| | BM | 35344.81 | 14740.4 | 0 | 34976.17 | 14963.4 | 0 |
| | EB | 35346.81 | 14742.4 | 0 | 34978.17 | 14965.4 | 0 |
| mammals | lambda | 13206.14 | 0 | 1 | 13057.58 | 0 | 1 |
| | OUr | 14050.39 | 844.2515 | 4.7E-184 | 13843.52 | 785.9418 | 2.2E-171 |
| | OUf | 14050.39 | 844.2515 | 4.7E-184 | 13843.52 | 785.9418 | 2.2E-171 |
| | BM | 20217.55 | 7011.412 | 0 | 19960.96 | 6903.375 | 0 |
| | EB | 20219.55 | 7013.412 | 0 | 19962.96 | 6905.375 | 0 |

**Supplementary Table 2**. Results of the phylogenetic regression of niche breadth (PC1 and PC2) and median aridity for all tetrapod clades. Note that the slopes (aridity estimates) are positive because low aridity index means high aridity, and therefore a positive slope here is indicating higher niche breadths in less arid species.

| | | PC1 breadth | | | | PC2 breadth | | | |
|---|---|---|---|---|---|---|---|---|---|
| | | Estimate | StdErr | t.value | p.value | Estimate | StdErr | t.value | p.value |
| amphibians | Intercept | 0.018314 | 0.417828 | 0.043831 | 0.96504 | 0.027387 | 0.42975 | 0.063727 | 0.94919 |
| | aridity | 0.304938 | 0.014565 | 20.93699 | 6.55E-94 | 0.327226 | 0.014509 | 22.55291 | 4.7E-108 |
| squamates | Intercept | -0.1914 | 0.504332 | -0.3795 | 0.704323 | -0.15648 | 0.49791 | -0.31427 | 0.75332 |
| | aridity | 0.381582 | 0.011683 | 32.6604 | 1.5E-221 | 0.418319 | 0.011659 | 35.87889 | 4.4E-264 |
| birds | Intercept | 0.077752 | 0.167503 | 0.464184 | 0.642529 | 0.105929 | 0.164755 | 0.64295 | 0.520276 |
| | aridity | 0.400299 | 0.011343 | 35.29089 | 2.3E-253 | 0.466004 | 0.010938 | 42.60462 | 0 |
| mammals | Intercept | 0.071417 | 0.468753 | 0.152355 | 0.878913 | 0.132309 | 0.43489 | 0.304234 | 0.760962 |
| | aridity | 0.281047 | 0.014451 | 19.44878 | 2.29E-81 | 0.336992 | 0.014183 | 23.76043 | 1.6E-118 |

**Supplementary Table 3**. Results of the phylogenetic regression between the residuals of niche breadth (PC1) on log-range size, and median aridity.

| | | Estimate | StdErr | t.value | p.value |
|---|---|---|---|---|---|
| amphibians | Intercept | -0.11945 | 0.362748 | -0.3293 | 0.741941 |
| | Estimate | 0.375915 | 0.013829 | 27.18226 | 2.7E-153 |
| squamates | Intercept | -0.09942 | 0.315031 | -0.3156 | 0.752313 |
| | Estimate | 0.424073 | 0.011042 | 38.40469 | 1.5E-299 |
| birds | Intercept | 0.011538 | 0.147541 | 0.078203 | 0.937668 |
| | Estimate | 0.465224 | 0.010964 | 42.43125 | 0 |
| mammals | Intercept | 0.078119 | 0.456427 | 0.171154 | 0.864109 |
| | Estimate | 0.403373 | 0.013856 | 29.11261 | 1.2E-172 |

**Supplementary Table 4**. Results of the multiple phylogenetic regressions between niche breadth (PC1 and PC2) and median aridity, controlling by log-range size by adding it as a predictor variable.

| | | PC1 breadth | | | | PC2 breadth | | | |
|---|---|---|---|---|---|---|---|---|---|
| | | Estimate | StdErr | t.value | p.value | Estimate | StdErr | t.value | p.value |
| amphibians | Intercept | -3.92245 | 0.309382 | -12.6784 | 2.35E-36 | -3.99181 | 0.296535 | -13.4615 | 1.06E-40 |
| | aridity | 0.309909 | 0.011523 | 26.89441 | 2.8E-150 | 0.332658 | 0.011259 | 29.54698 | 6.4E-179 |
| | log(range size) | 0.173007 | 0.002988 | 57.9097 | 0 | 0.176406 | 0.002922 | 60.3735 | 0 |
| squamates | Intercept | -3.62247 | 0.269716 | -13.4307 | 9.61E-41 | -3.67569 | 0.247102 | -14.8752 | 1.78E-49 |
| | aridity | 0.344228 | 0.00932 | 36.93391 | 1.1E-278 | 0.383519 | 0.009124 | 42.03196 | 0 |
| | log(range size) | 0.153418 | 0.00204 | 75.22016 | 0 | 0.156358 | 0.002008 | 77.87655 | 0 |
| birds | Intercept | -4.39231 | 0.152759 | -28.7532 | 6E-173 | -4.04647 | 0.158239 | -25.5719 | 1.3E-138 |
| | aridity | 0.443016 | 0.009837 | 45.03566 | 0 | 0.504366 | 0.009634 | 52.35115 | 0 |
| | log(range size) | 0.166031 | 0.003353 | 49.51084 | 0 | 0.154094 | 0.003276 | 47.03513 | 0 |
| mammals | Intercept | -4.41099 | 0.397868 | -11.0866 | 3.03E-28 | -4.32192 | 0.343539 | -12.5806 | 9E-36 |
| | aridity | 0.351797 | 0.012017 | 29.27486 | 2E-174 | 0.410143 | 0.011687 | 35.09345 | 3.8E-242 |
| | log(range size) | 0.176629 | 0.003597 | 49.11142 | 0 | 0.175415 | 0.003527 | 49.73179 | 0 |

**Supplementary Table 5**. Phylogenetic regression between niche breadth (PC1 and PC2) and the percentage of arid cells in species' distribution ranges.

| | | PC1 breadth | | | | PC2 breadth | | | |
|---|---|---|---|---|---|---|---|---|---|
| | | Estimate | StdErr | t.value | p.value | Estimate | StdErr | t.value | p.value |
| amphibians | Intercept | -0.00603 | 0.462216 | -0.01304 | 0.989594 | -0.00312 | 0.484043 | -0.00645 | 0.994855 |
| | % arid cells | -0.43068 | 0.080952 | -5.32018 | 1.08E-07 | -0.39278 | 0.080959 | -4.85165 | 1.26E-06 |
| squamates | Intercept | -0.05939 | 0.324455 | -0.18305 | 0.854765 | -0.04415 | 0.324357 | -0.13613 | 0.891721 |
| | % arid cells | -0.6608 | 0.031017 | -21.3046 | 2.3E-98 | -0.68085 | 0.031259 | -21.7811 | 1.3E-102 |
| birds | Intercept | 0.231782 | 0.177801 | 1.303602 | 0.192408 | 0.247722 | 0.199293 | 1.243007 | 0.213902 |
| | % arid cells | -1.13648 | 0.044457 | -25.5635 | 1.6E-138 | -1.06561 | 0.04452 | -23.9355 | 2.9E-122 |
| mammals | Intercept | 0.262582 | 0.517609 | 0.507297 | 0.611968 | 0.331995 | 0.511713 | 0.648791 | 0.516502 |
| | % arid cells | -0.73154 | 0.048159 | -15.1901 | 5.14E-51 | -0.70012 | 0.048284 | -14.5 | 9.93E-47 |

**Supplementary Table 6**. Results of the phylogenetic ANOVAs comparing the niche breadth between arid and non-arid species.

| | | Df | SS | MS | Rsq | F | Z | Pr(>F) |
|---|---|---|---|---|---|---|---|---|
| amphibians | arid | 1 | 11.90312 | 11.90312 | 0.001522 | 8.889462 | 2.359074 | 0.007 |
| | Residuals | 5830 | 7806.455 | 1.339015 | 0.998478 | | | |
| | Total | 5831 | 7818.358 | | | | | |
| squamates | arid | 1 | 316.2782 | 316.2782 | 0.020904 | 197.8282 | 7.085863 | 0.001 |
| | Residuals | 9266 | 14814.04 | 1.598752 | 0.979096 | | | |
| | Total | 9267 | 15130.32 | | | | | |
| birds | arid | 1 | 1077.976 | 1077.976 | 0.066557 | 560.0082 | 7.595356 | 0.001 |
| | Residuals | 7854 | 15118.39 | 1.924929 | 0.933443 | | | |
| | Total | 7855 | 16196.37 | | | | | |
| mammals | arid | 1 | 308.446 | 308.446 | 0.011785 | 61.55885 | 4.660999 | 0.001 |
| | Residuals | 5162 | 25864.65 | 5.010588 | 0.988215 | | | |
| | Total | 5163 | 26173.1 | | | | | |
