## Appendix 1 for "Aridity Drives Climatic Specialization and Phylogenetic Clustering in Terrestrial Vertebrates"

### Cleaning bird distribution files

2025-01-17

#### Introduction

This report filters bird distribution data to include only breeding areas and processes polygon geometries.

---

#### Load Required Packages

```
# Load packages
packages <- c('tidyverse', 'sf', 'rgeos', 'rgdal', 'terra', 'phytools', 'geiger')
easypackages::libraries(packages)

sf_use_s2(FALSE)
```

---

#### Data Import

##### BOTW Data

```
# Import data
botw_file <- ''
botw_folder <- ''

botw_layers <- st_layers(botw_folder)
botw <- st_read(dsn = botw_folder, layer = 'All_Species', quiet = FALSE)
botw_table <- st_read(dsn = botw_folder, layer = 2)

head(botw)
```

---

#### Data Cleaning

##### Filtering Data

```r
# Filter the data to include only:
# - Extant species in the column 'presence' (number 1).
# - Native species in the column 'origin' (number 1).
# - Resident throughout the year (1) or breeding (2) in the column 'seasonal'.
botw_clean <- botw %>%
  filter(presence == 1) %>%
  filter(origin == 1) %>%
  filter(seasonal %in% c(1, 2))

nrow(botw)
nrow(botw_clean)

saveRDS(botw_clean, 'data/distribution/botw_clean.rds')
```

---

### Validating Polygons

#### Checking and Fixing Geometries

```r
# Create list of polygons
botw_clean$id <- 1:nrow(botw_clean)

# Make a list of separate species range polygons
spList <- vector('list', nrow(botw_clean))
names(spList) <- botw_clean$id

for (i in 1:length(spList)) {
  spList[[i]] <- botw_clean[i,]
}

problematic_polygons <- c()
for (i in 1:length(spList)) {
  print(paste0('polygon ', i, ' / ', length(spList), ' --- ', botw_clean$sci_name[i]))
  if (st_geometry_type(spList[[i]]) == 'MULTISURFACE'){
    message('MULTISURFACE polygon', i)
    problematic_polygons <- c(problematic_polygons, i)
  } else {
    if (!any(st_is_valid(spList[[i]]))) {
      message('\trepairing poly ', i)
      spList[[i]] <- st_make_valid(spList[[i]])
      if (!any(st_is_valid(spList[[i]]))) {
        message('\t\tstill broken...')
      }
    }
  }
}

# Save results
saveRDS(spList, 'output/spList_polygons_birds.rds')
saveRDS(problematic_polygons, 'output/problematic_polygons_birds.rds')
```

---

### Grouping Data by Species

#### Final Processing

```
# Group by species
spList <- readRDS('output/spList_polygons_birds.rds')
problematic_polygons <- readRDS('output/problematic_polygons_birds.rds')

botw_clean_valid <- data.table::rbindlist(spList, use.names = TRUE)
botw_clean_valid <- botw_clean_valid[-c(problematic_polygons),]

botw_clean_valid_grouped <- botw_clean_valid %>%
  group_by(sci_name) %>%
  summarize(Shape = st_union(Shape)) %>%
  st_as_sf()

saveRDS(botw_clean_valid_grouped, 'output/botw_clean_valid_grouped.rds')
```

---

### Conclusion

This workflow filtered bird distribution data, addressed polygon validity issues, and grouped distribution ranges by species. The final output is saved for further analyses.
