## Supplementary figures and images for "Aridity Drives Climatic Specialization and Phylogenetic Clustering in Terrestrial Vertebrates"

### Appendix 2

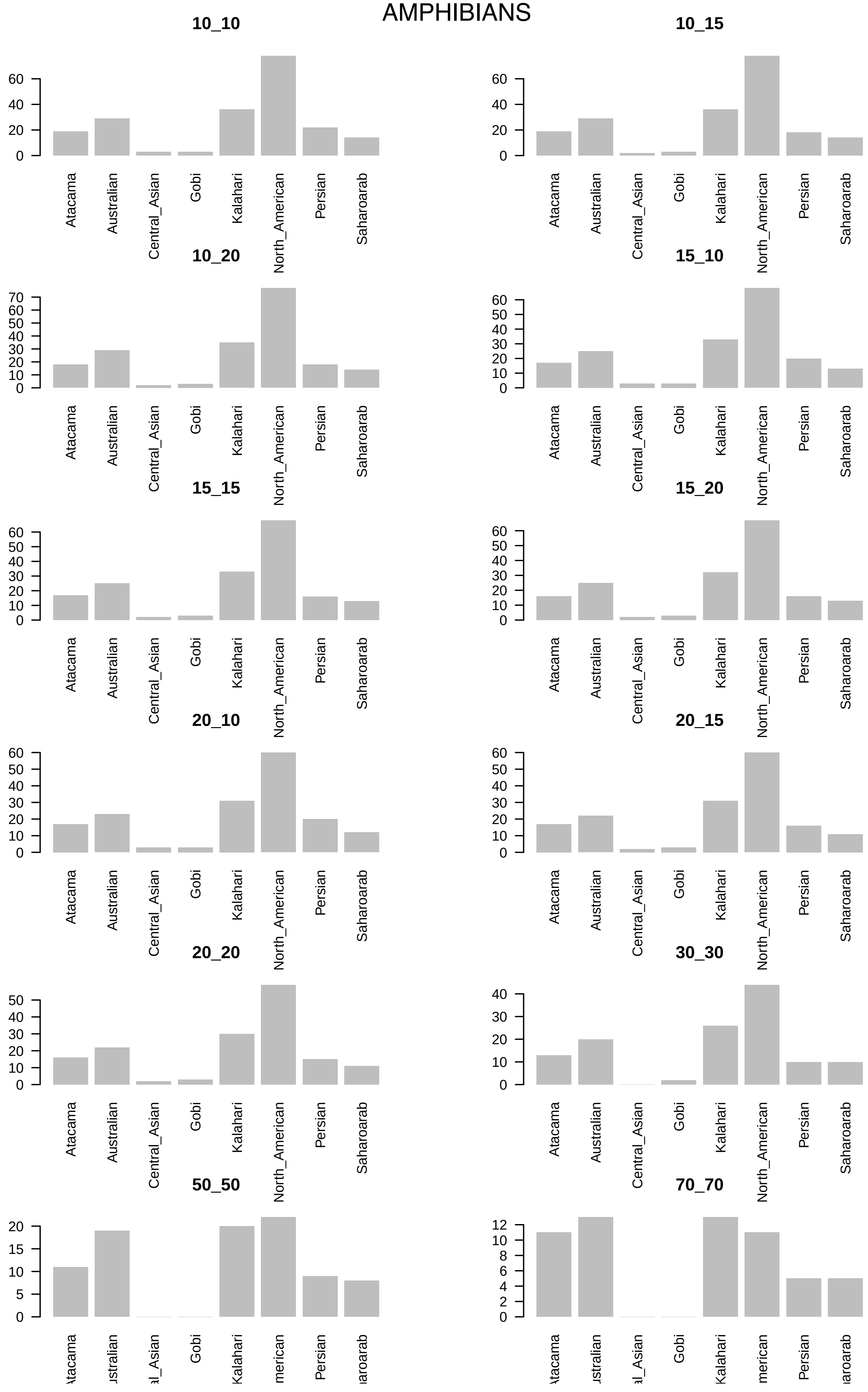

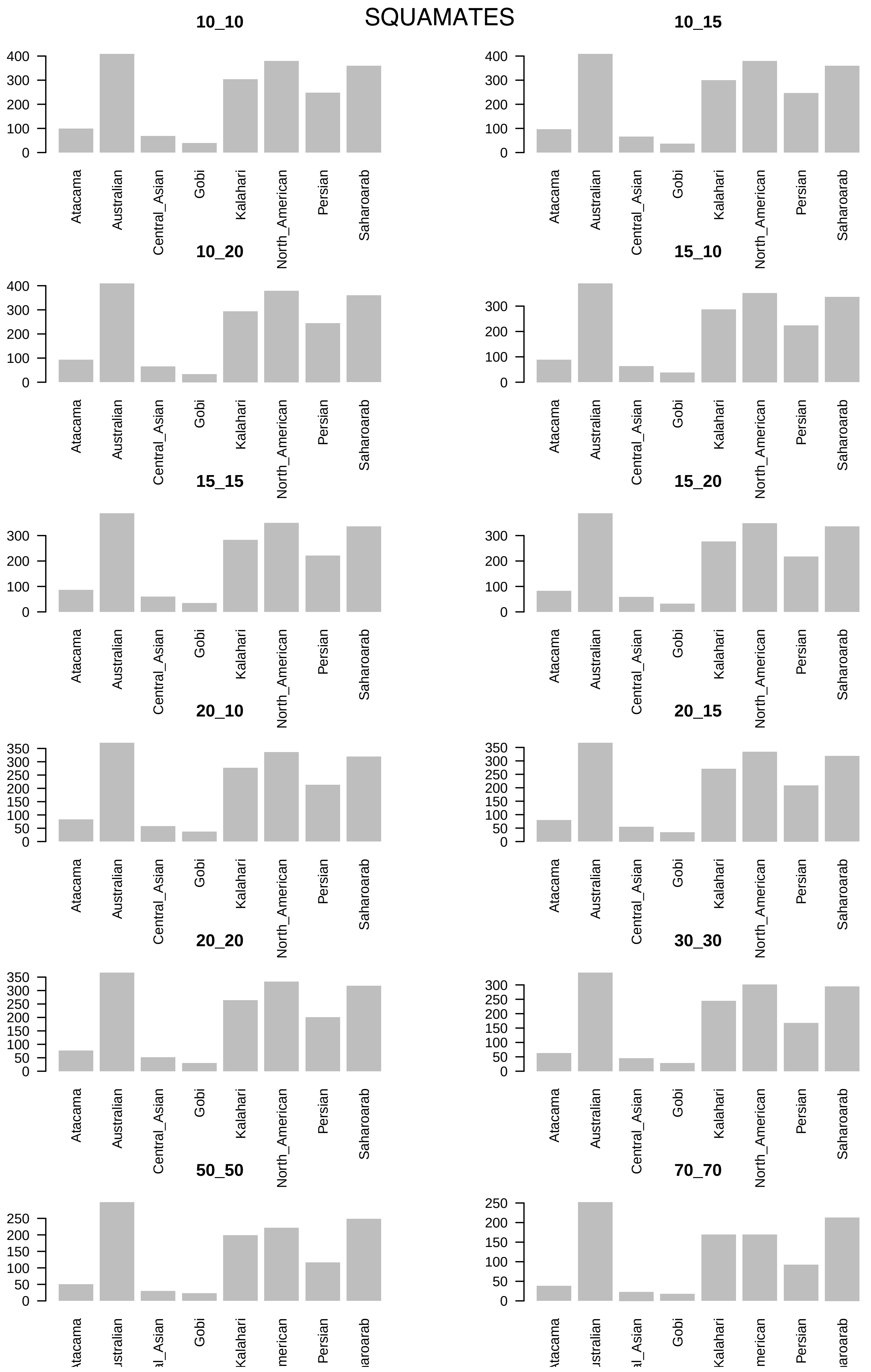

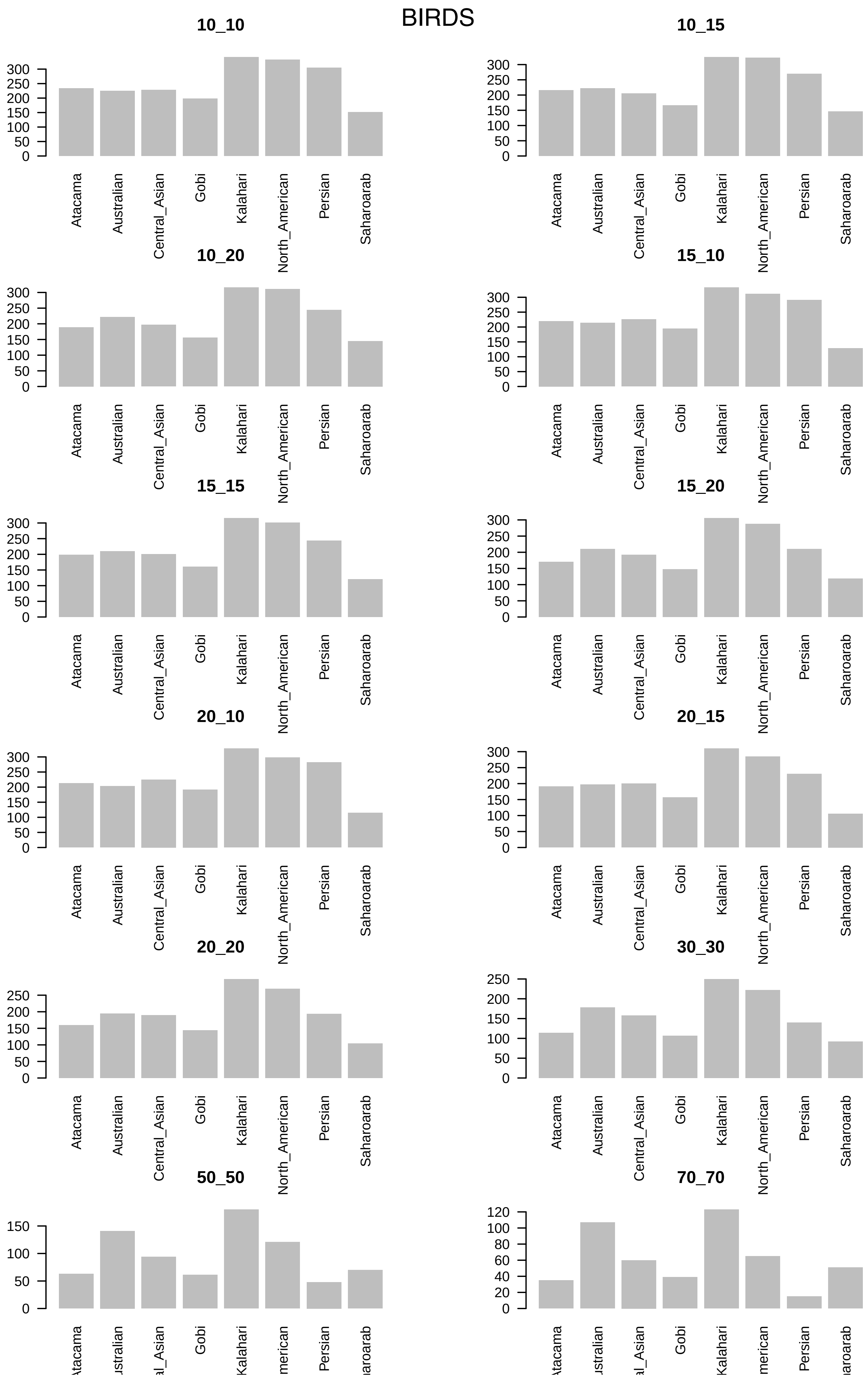

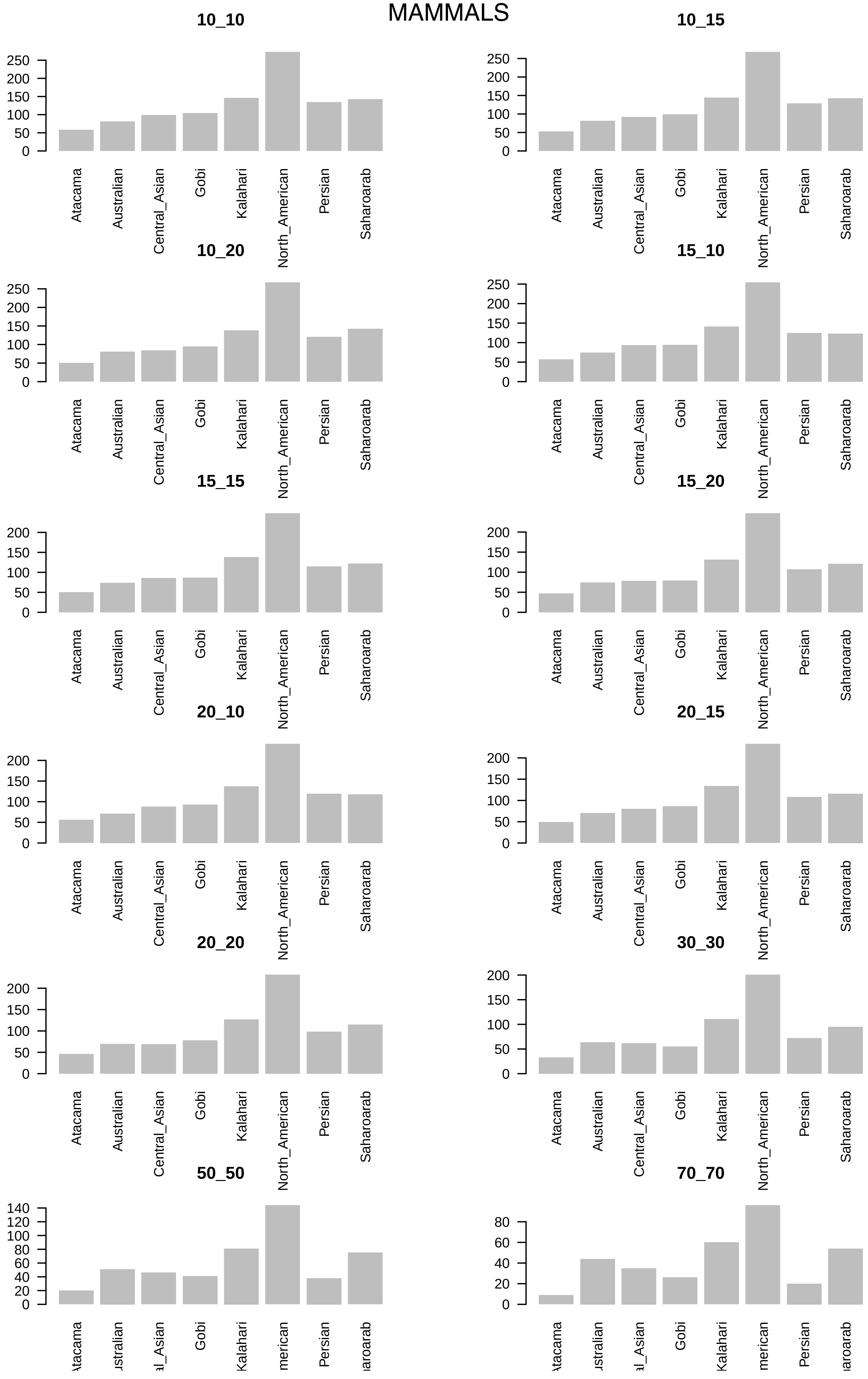
